# Design rules for efficient endosomal escape

**DOI:** 10.1101/2023.11.03.565388

**Authors:** Madeline Zoltek, Angel Vázquez, Xizi Zhang, Neville Dadina, Lauren Lesiak, Alanna Schepartz

**Author notes:** Alanna Schepartz, **Email:**. **Author Contributions:** Study conception and design: M.Z. and A.S.; preparation of materials: M.Z., A.V., X.Z. and L.L.; data collection: M.Z., A.V., X.Z., and N.D.; analysis and interpretation of results: M.Z., A.V., and N.D.; and manuscript preparation: M.Z. and A.S. **Competing Interest Statement:** None.

## Abstract

The inefficient translocation of proteins across biological membranes limits their application as therapeutic compounds and research tools. In most cases, translocation involves two steps: uptake into the endocytic pathway and endosomal escape. Certain charged or amphiphilic molecules promote protein uptake but few enable efficient endosomal escape. One exception is ZF5.3, a mini-protein that exploits natural endosomal maturation machinery to translocate across endosomal membranes. Although certain ZF5.3-protein conjugates are delivered efficiently into the cytosol or nucleus, overall delivery efficiency varies widely with no obvious design rules. Here we evaluate the role of protein size and thermal stability in the ability to efficiently escape endosomes when attached to ZF5.3. Using fluorescence correlation spectroscopy, a singlemolecule technique that provides a precise measure of intra-cytosolic protein concentration, we demonstrate that delivery efficiency depends on both size and the ease with which a protein unfolds. Regardless of size and pI, low-Tm cargos of ZF5.3 (including intrinsically disordered domains) bias its endosomal escape route toward a high-efficiency pathway that requires the homotypic fusion and protein sorting (HOPS) complex. Small protein domains are delivered with moderate efficiency through the same HOPS portal even if the Tm is high. These findings imply a novel protein- and/or lipid-dependent pathway out of endosomes that is exploited by ZF5.3 and provide clear guidance for the selection or design of optimally deliverable therapeutic cargo.

**Significance Statement:** The results described in this paper provide new insights into how protein delivery works and how it can be best utilized in the future. Although intracellular protein delivery has been studied for decades, this paper describes the first interrogation of why certain protein cargos are privileged for efficient endosomal escape. These results represent a fundamental advance in the long-awaited goal of efficient protein delivery and provide design rules to overcome one of the most significant challenges for the future of biotechnology.

## Introduction

Protein- and nucleic acid-derived biologics are a rapidly expanding sector of modern drug development. When compared to small molecules, biologics can improve target specificity, inhibit or activate recalcitrant targets, replace missing or malfunctioning enzymes, and deliver gene editing or protein-editing machineries (1). Direct protein delivery is simpler than lipid nanoparticle or viral vector delivery strategies (2) and provides fine-tuned control over dosage and intracellular lifetime. Despite this potential, there is not a single approved protein therapeutic that operates in the cytosol or nucleus. The problem is inefficient endosomal escape. Decades of research dedicated to improving endosomal escape of proteins delivered via the endosomal pathway has yielded many molecules that stimulate endocytic uptake, but almost none that escape endosomes and avoid a degradative fate.

One molecule that has shown significant promise with regard to endosomal escape is ZF5.3, a 27-aa mini-protein that exploits the HOPS complex, a natural and ubiquitous component of the endosomal maturation machinery (3–6), to guide certain proteins into the cytosol and nucleus with exceptional efficiency (6–9). A conjugate of ZF5.3 and the transcriptional repressor MeCP2 (implicated in Rett Syndrome) reaches the nucleus of mammalian cells with an efficiency of >80% (defined as nuclear concentration divided by treatment concentration) while retaining its native binding partners and function (6). Notably, delivery of ZF5.3-MeCP2 is substantially more efficient than other ZF5.3-protein conjugates (7, 8) and, to our knowledge, any other reported nucleic acid or protein biologic that escapes the endocytic pathway. Precisely which attributes of ZF5.3-MeCP2 enabled such efficient endosomal escape, and whether these attributes could be generalized, however, remained unclear. Equally unclear was how an endosomal maturation machinery that catalyzes vesicle fusion from the cytosol (10) communicates across a membrane barrier with ZF5.3-protein cargo located within the endosomal lumen.

Endosomal escape of a biologic inherently requires the energetically unfavorable translocation of a hydrophilic molecule across a hydrophobic membrane. Nature overcomes the challenges of protein translocation through two fundamentally distinct mechanisms: one that requires unfolding of the protein being transported (e.g. via Sec-translocases (11, 12) or mitochondrial import pathways (13–15)), and one that accommodates the globular fold of the protein in transit (e.g. during peroxisome entry (16) or unconventional protein secretion (17, 18)). Regardless of the cellular machinery required, given that the structure of MeCP2 is up to 60% disordered (6, 19), we hypothesized that intrinsic disorder could favor endosomal escape through a pathway that demands protein unfolding.

Here we test this hypothesis, and discover that the ability to unfold is a key determinant in how well ZF5.3 guides a protein into the cytosol in a HOPS-dependent manner. Proteins that are intrinsically disordered or unfold at physiological temperatures are delivered into the cytosol by ZF5.3 with high efficiency and in a HOPS-dependent manner; proteins with greater stability can be delivered with modest efficiency and in a HOPS-dependent manner if the domain is sufficiently compact. Super-resolution microscopy images of endolysosomes in ZF5.3-treated cells provide evidence for distinct condensed sub-populations that associate with the limiting membrane. Our data support a model in which intrinsically disordered proteins or those that unfold readily are privileged with respect to efficient endosomal escape via a HOPS-dependent portal. We anticipate that these design rules will constitute a useful filter in the further development of direct protein delivery strategies and provide new insights into how proteins, natural or designed, circumnavigate biological membranes.

## Results

To establish whether unfolding plays a role in ZF5.3-mediated endosomal escape, we built on classic work of Eilers & Schatz, who almost forty years ago utilized the ligand-dependent stability of dihydrofolate reductase (DHFR) to study protein import into mitochondria (13). The thermal stability of DHFR (T_m_) increases by approximately 15 °C upon the binding of ligands such as methotrexate (MTX) or trimethoprim (13). Indeed, the effect of MTX or trimethoprim on protein import and export established a role for protein unfolding during chaperone-mediated lysosomal import mediated by heat shock family molecular chaperones (20, 21), protein translocation across the *E. coli* plasma membrane mediated by the Sec-translocase (11, 12), endoplasmic reticulum retrotranslocation (22), and cytosolic delivery of toxins such as ricin and diphtheria (23–25).

We purified samples of DHFR and ZF5.3-DHFR from *E. coli* and confirmed their identities using SDS-PAGE and LC/MS, respectively (*SI Appendix*, Fig. S1a-c). The presence of ZF5.3 at the N-terminus of DHFR has little or no effect on overall protein secondary structure or catalytic activity (*SI Appendix*, Fig. S1d,e). With these materials in hand, we established baseline values for the cytosolic delivery of DHFR and ZF5.3-DHFR using rhodamine-tagged variants (DHFR^Rho^ and ZF5.3-DHFR^Rho^) prepared using sortase, as described previously *(SI Appendix*, Fig. S1a-c) (6–8). We incubated human osteosarcoma (Saos-2) cells with 100 - 1000 nM DHFR^Rho^ or ZF5.3-DHFR^Rho^ for 1 h, washed and trypsin-treated the cells to remove surface-bound material, and visualized the cells using confocal microscopy, flow cytometry (FC), and fluorescence correlation spectroscopy (FCS) (Fig. 1 and *SI Appendix*, Fig. S2,3). Confocal microscopy and FC revealed that cells treated with ZF5.3-DHFR^Rho^ showed substantially higher total intracellular fluorescence than those treated with DHFR^Rho^ at all treatment concentrations and time points. The overall uptake of DHFR^Rho^ and ZF5.3-DHFR^Rho^ revealed by confocal microscopy (Fig. 1a and *SI Appendix*, Fig. S2) and FC (Fig. 1b) was dose-dependent; the total uptake of ZF5.3-DHFR^Rho^ was significantly higher than that of DHFR^Rho^, especially at treatment concentrations of 500 nM (15.5-fold increase) and 1 µM (30.6-fold increase). These increases in total uptake due to fusion to ZF5.3 are in line with values measured for other ZF5.3-protein conjugates (7, 8). No increase in uptake was observed when cells were treated with a 1:1 mixture of ZF5.3 and DHFR^Rho^ (Fig. 1b), confirming that a covalent linkage is required for enhanced delivery (8).

**Fig 1.**
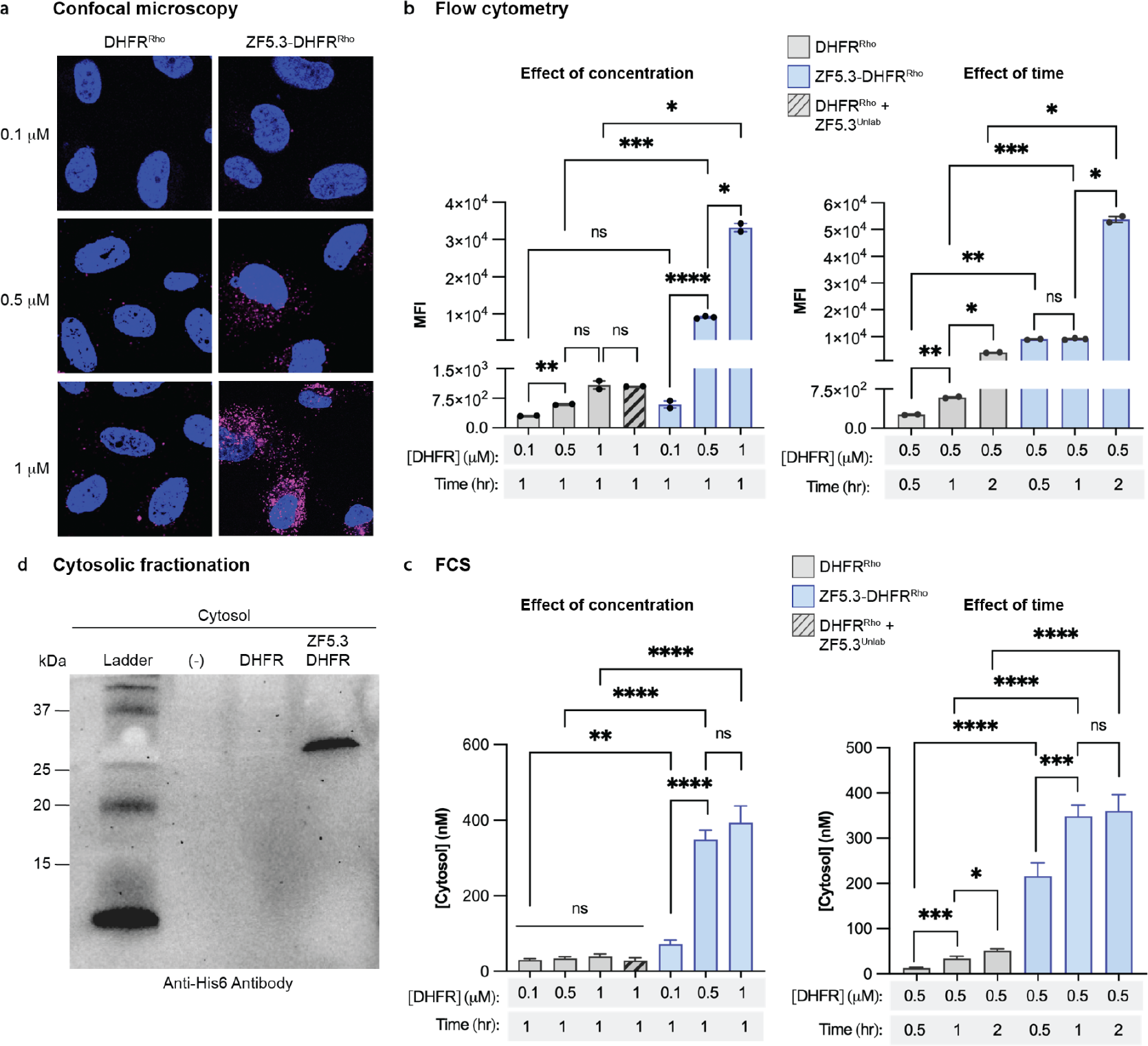
DHFR reaches the cytosol efficiently when covalently fused to ZF5.3. a, 2-D confocal microscopy images of Saos-2 cells incubated with the indicated concentration of DHFRRho or ZF5.3-DHFRRho as described in Online Methods. Plots showing b, flow cytometry analysis of total cellular uptake or c, FCS analysis of cytosolic concentration of DHFRRho, ZF5.3-DHFRRho, or a 1:1 mixture of ZF5.3 and DHFRRho after the indicated treatment concentration and incubation time; see Online Methods for detailed procedure. Flow cytometry values are provided as Median Fluorescence Intensity (MFI) for the lissamine rhodamine B channel, n = 20000 in total per condition containing at least 2 biological replicates each (mean ± SEM). FCS values provided in nM, n>20 for each FCS condition with two biological replicates each (mean ± SEM). Statistical significance comparing the given concentrations was assessed using the Brown-Forsythe and Welch one-way analysis of variance (ANOVA) followed by an unpaired t-test with Welch’s correction. ****p ≤ 0.0001, ***p ≤ 0.001, **p ≤ 0.01, *p ≤ 0.05. d, Western blot analysis of fractionated cytosol from Saos-2 cells treated with either DMEM media alone (-), DHFR, or ZF5.3-DHFR at 0.5 µM for 1 h. The presence of intact DHFR or ZF5.3-DHFR was assessed using an anti-His6 antibody. The gel results shown are representative of two biological replicates.

Although endocytic uptake is the first step along the pathway to the cytosol, the key determinant of delivery efficiency is endosomal escape – the fractional concentration of intact protein that reaches the cytosol. Two challenges have thwarted attempts to improve cytosolic delivery. The first is the absence of tools to accurately quantify delivery efficiency, and the second is the difficulty in establishing whether the delivered material is intact (or not) and thus capable of function. We used live cell FCS (26) to establish delivery efficiency (27) by quantifying the concentration of DHFR^Rho^ and ZF5.3-DHFR^Rho^ that reached the cytosol of Saos-2 cells. Unlike flow cytometry, FCS provides both the concentration and the diffusion time of a fluorescent molecule within a subcellular compartment, such as the cytosol or nucleus (27–29). The former value provides an accurate measure of delivery efficiency, while the latter, when combined with careful biochemistry, establishes whether the fluorescent material is intact (27, 30).

### ZF5.3-DHFR^Rho^ trafficks efficiently into the Saos-2 cytosol

Examination of treated Saos-2 cells using FCS revealed substantial differences in the efficiencies with which DHFR^Rho^ and ZF5.3-DHFR^Rho^ reached the cytosol. Cells treated with DHFR^Rho^ showed little trafficking of this material to the cytosol at any concentration studied (Fig. 1c and *SI Appendix*, Fig. S3). At the highest treatment concentration (1 µM) the measured cytosolic concentration of DHFR^Rho^ was 39 nM, a delivery efficiency of only 3.9%. By contrast, ZF5.3-DHFR^Rho^ reached the cytosol efficiently and in a dose-dependent manner, establishing average concentrations of 72, 350, and 470 nM when cells were treated with 100, 500, and 1000 nM ZF5.3-DHFR^Rho^, respectively, for 1 h (Fig. 1c). These values correspond to delivery efficiencies between 47 and 72%, up to 10-fold higher than those measured for DHFR^Rho^. Notably, at a fixed treatment concentration of 500 nM ZF5.3-DHFR^Rho^, additional incubation time (up to 2 hr) improves total uptake but does not substantially increase the fraction that reaches the cytosol (Fig. 1b,c). These data suggest that ZF5.3-DHFR^Rho^ follows a saturable pathway to escape from endosomes, and that endosomal escape (as opposed to an earlier endocytic event) kinetically limits delivery to the cytosol. When stringently isolated from the cytosol of treated cells, ZF5.3-DHFR was recovered fully intact with no evidence of either degradation or endosomal contamination (Fig. 1d and *SI Appendix*, Fig. S4). Co-administration of ZF5.3 did not improve the cytosolic delivery of DHFR^Rho^, confirming that efficient delivery demands a covalent linkage to ZF5.3^8^ (Fig 1c). Thus, the presence of ZF5.3 at the N-terminus of DHFR^Rho^ improved delivery to the cytosol by up to 12-fold. The cytosolic delivery of ZF5.3-DHFR^Rho^ is more efficient than nearly all other proteins delivered by ZF5.3 previously (7, 8), and though it is not intrinsically disordered, the translocation efficiency of ZF5.3-DHFR^Rho^ into the cytosol mirrors that of ZF5.3-MeCP2 (6).

### Delivery of DHFR by ZF5.3 is inhibited by equimolar MTX

Next, to interrogate the role of protein folding in cytosolic delivery mediated by ZF5.3, we determined the impact of the DHFR-selective inhibitor methotrexate (MTX, Fig. 2a) on the cytosolic delivery efficiencies of DHFR^Rho^ and ZF5.3-DHFR^Rho^. MTX binds DHFR with sub-nanomolar affinity (K_D_ ≈ 10^-10^ M) (31) and potently inhibits enzyme activity (32) (*SI Appendix*, Fig. S1e). Temperature-dependent circular dichroism (CD) spectroscopy established that the apparent thermal stabilities (*T_m_) of DHFR and ZF5.3-DHFR increased by approximately 15 degrees in the presence of 1 equivalent MTX. For DHFR, the *T_m_ measured in the absence of MTX was 44.5°C, in line with previous measurements (33), and increased by 16.6 °C in the presence of 1 equivalent MTX. For ZF5.3-DHFR, the *T_m_ in the absence of MTX was 32.7°C and the corresponding increase was 17.3°C (Fig. 2b).

**Fig. 2.**
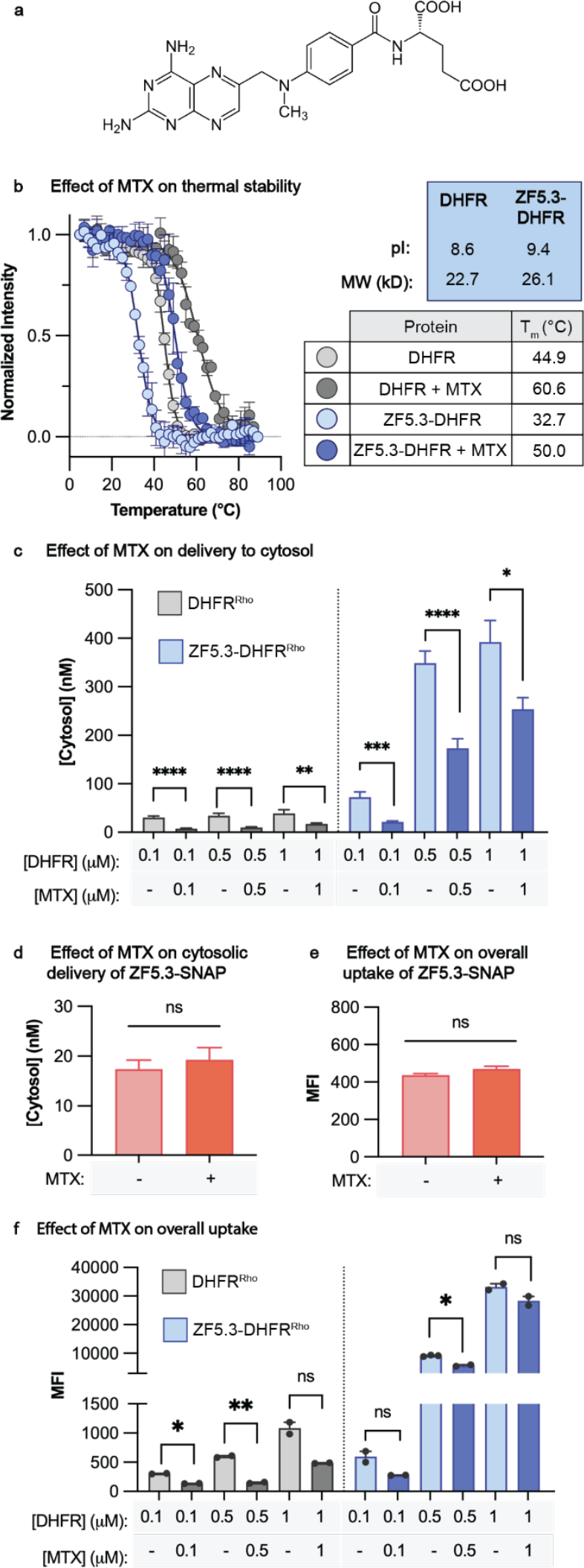
Delivery of DHFR by ZF5.3 is inhibited by equimolar MTX. **a**, Chemical structure of methotrexate (MTX). **b**, Plots illustrating the temperature-dependent loss in circular dichroism (CD) signal at 210 nm for DHFR and ZF5.3-DHFR (20 µM protein in 25 mM Tris, 150 mM KCl, 1 mM TCEP, pH 7.2) in the presence or absence of 1 equivalent MTX. For each melt, the temperature was increased in 2° increments between 5 and 90°C and the ellipticity at 210 nm was fitted using a Boltzmann sigmoidal nonlinear regression. The melts were irreversible and therefore we report the midpoint value of these fitted curves as apparent T_m_ values (*T_m_). The data shown include two biological replicates. **c-f**, Plots illustrating the effect of MTX on the cytosolic delivery (c, d) and overall uptake (e, f) of DHFR^Rho^, ZF5.3-DHFR^Rho^, or ZF5.3-SNAP^Rho^. In all cases, the proteins were pre-incubated with 1 equivalent MTX for 30 minutes before treatment at 500 nM for 1 h (DHFR^Rho^ and ZF5.3-DHFR^Rho^) or 1000 nM for 30 min (ZF5.3-SNAP^Rho^). Cells were then trypsinized and analyzed by flow cytometry or FCS as described in Online Methods. Flow cytometry values are provided as Median Fluorescence Intensity for the lissamine rhodamine B channel (MFI), n = 20000 per condition in total with at least 2 biological replicates each (mean ± SEM). FCS values provided in nM, n>20 for each FCS condition comprising two biological replicates each (mean ± SEM). Statistical significance comparing the given concentrations was assessed using the Brown-Forsythe and Welch one-way analysis of variance (ANOVA) followed by an unpaired t-test with Welch’s correction. ****p ≤ 0.0001, ***p ≤ 0.001, **p ≤ 0.01, *p ≤ 0.05.

Samples of DHFR^Rho^ and ZF5.3-DHFR^Rho^ at concentrations from 100 - 1000 nM were preincubated with 1 equivalent MTX for 30 minutes, added to Saos-2 cells, and incubated for 1 h as described previously. Under all conditions, the presence of 1 equivalent MTX substantially decreased the fraction of ZF5.3-DHFR^Rho^ that reached the cytosol (Fig. 2c). The effect of MTX was inversely related to ZF5.3-DHFR^Rho^ concentration, with reductions of 70.4%, 50.4%, and 42.8% at incubation concentrations of 100, 500, and 1000 nM, respectively (Fig. 2c). Notably, MTX also decreased the concentration of DHFR^Rho^ that reached the cytosol by comparable amounts, but had no effect on the cytosolic delivery of ZF5.3-SNAP^Rho^, an unrelated protein (Fig. 2d).

To evaluate the extent to which MTX affected cytosolic delivery by inhibiting overall uptake of ZF5.3-DHFR^Rho^, we also evaluated treated cells using flow cytometry (Fig. 2e). These results indicate that MTX has different effects on the overall uptake of DHFR^Rho^ and ZF5.3-DHFR^Rho^. Although one equivalent of MTX substantially decreased the overall uptake of DHFR^Rho^ by between 55 and 75% at all treatment concentrations, there was little or no effect of MTX on the overall uptake of ZF5.3-DHFR^Rho^ at treatment concentrations of 500 and 1000 nM. MTX had no effect on the overall uptake of the unrelated protein ZF5.3-SNAP^Rho^ (Fig. 2e). The observation that MTX has a substantial effect on delivery of ZF5.3-DHFR^Rho^ to the cytosol but little or no effect on overall uptake implies that unfolding plays a significant role in one or more of the steps that guides ZF5.3-DHFR^Rho^ out of the endocytic pathway and into the cytosol. For this reason, the relatively low thermostability (T_m_ = 32.7°C) of ZF5.3-DHFR likely contributes to its highly efficient endosomal escape. These data also suggest that endosomal uptake and escape of DHFR^Rho^ and ZF5.3-DHFR^Rho^ proceed using fundamentally different molecular machinery or pathways, but only the pathway accessed by ZF5.3-DHFR results in efficient cytosolic delivery.

### Unfolding of cargo is a general requirement for high-efficiency ZF5.3 delivery

Although one equivalent MTX inhibits the fraction of ZF5.3-DHFR^Rho^ that reaches the cytosol (Fig. 2c), the inhibition is partial, not complete. We reasoned that this finding might be due to the loss of MTX from ZF5.3-DHFR^Rho^ before the complex reaches the late endosomal compartment from which escape occurs, especially as the compartments become progressively more acidic. To more directly evaluate the role of unfolding in endosomal escape, we turned to three known SNAP-tag variants that differ by only a few amino acid substitutions but nonetheless show distinctly different thermal stabilities (34–37). These variants, all intermediates generated along the directed evolution pathway between human O^6^-alkylguanine-DNA alkyltransferase and commercially available SNAP-tag, display thermal stabilities between 35 - 51 °C but with nearly indistinguishable molecular weights and isoelectric points of 8.7 ± 0.1 (Fig. 3a and *SI Appendix*, Fig. S5a) (34). Each SNAP-tag variant was conjugated to the C-terminus of ZF5.3 and tagged with rhodamine upon reaction with benzylguanine-modified lissamine rhodamine B (BG-Rho) (*SI Appendix*, Fig. S5b,c). Temperature-dependent CD studies confirmed the previously reported thermal stabilities; once again, the presence of ZF5.3 had a modest destabilizing effect on the *T_m_ but little or no effect on overall secondary structure (Fig. 3b and *SI Appendix*, Fig. S5d).

**Fig. 3:**
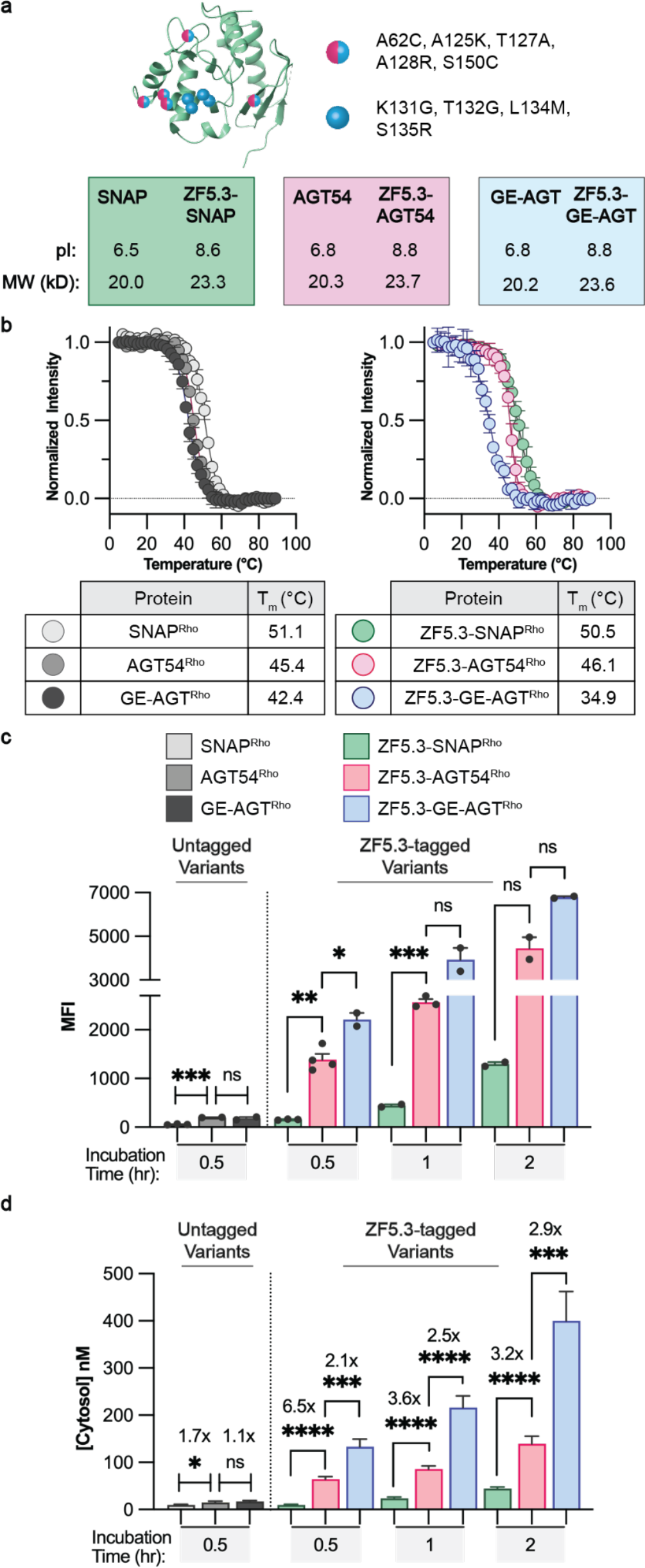
Destabilization of SNAP-tag enhances intracellular delivery only when conjugated to ZF5.3. **a**, Structure of previously published SNAP-tag (PDB: 6Y8P) with destabilizing mutations marked by magenta and blue circles (for residues found in both AGT54 and GE-AGT) or blue circles (for residues found only in GE-AGT). Isoelectric point (pI) and molecular weight (in kilo-daltons, kD) are increased with the addition of ZF5.3 but remain highly similar among all variants. **b**, Plots illustrating the temperature dependence of the 222 nm CD signal of GE-AGT^Rho^, AGT54^Rho^, and SNAP^Rho^ alongside the corresponding ZF5.3 conjugates. Each protein at 20 µM was measured in 20 mM Tris, 150 mM NaCl, pH 7.5. For each melt, the temperature was increased in 2° increments between 5 and 90°C and the ellipticity at 222 nm was fitted using a Boltzmann sigmoidal nonlinear regression to obtain *T_m_ values. The data shown include two biological replicates. **c, d**,Total cellular uptake (top) and cytosolic concentration (bottom) of the untagged or ZF5.3-tagged SNAP^Rho^ variants were analyzed by flow cytometry and FCS. Saos-2 cells were incubated with 1 µM of the given protein for 30 min, 1 h, or 2 h before the cellular workup and measurements were performed as described previously. Flow cytometry values are provided as Median Fluorescence Intensity for the lissamine rhodamine B channel, n = 20000 per condition in total with two biological replicates each (mean ± SEM). FCS values are provided in nM, n>20 for each FCS condition with two biological replicates each (mean ± SEM). Statistical significance was assessed using the Brown-Forsythe and Welch one-way analysis of variance (ANOVA) followed by an unpaired t-test with Welch’s correction. ****p ≤ 0.0001, ***p ≤ 0.001, **p ≤ 0.01, *p ≤ 0.05.

Saos-2 cells were treated with each SNAP^Rho^ variant (1 µM) for 0.5 - 2 h and evaluated using confocal microscopy, flow cytometry, and FCS as described previously (Fig. 3c,d and *SI Appendix*, Fig. S6 and 7). The most stable variant (ZF5.3-SNAP^Rho^, *T_m_ = 51°C) showed minimal uptake (Fig. 3c) and poor trafficking to the cytosol (Fig. 3d) regardless of incubation time, in line with results described previously for a closely related variant (7). The less thermostable proteins, ZF5.3-GE-AGT^Rho^ (*T_m_ = 35°C) and ZF5.3-AGT54^Rho^ (*T_m_ = 46°C), were taken up with higher efficiency but not equally when evaluated by flow cytometry, with uptake increasing after longer incubation times (Fig. 3c). Given the roughly equal surface charges of SNAP, AGT54, and GE-AGT, it is interesting to note that decreased thermal stability seems to improve overall ZF5.3-mediated cellular uptake.

Notably, the three ZF5.3-SNAP^Rho^ variants trafficked to the cytosol with different efficiencies, and in a manner that correlated directly with *T_m_ (Fig. 3d and *SI Appendix*, Fig. S6). At all incubation times, ZF5.3-GE-AGT^Rho^, with the lowest *T_m_ (35°C), reached the cytosol between 2.1 - 2.9 -fold more efficiently than mid-*T_m_ ZF5.3-AGT54^Rho^, which in turn reached the cytosol 3.2 - 6.5-fold more efficiently than high-*T_m_ ZF5.3-SNAP^Rho^. At its maximum, the least thermostable variant ZF5.3-GE-AGT^Rho^ reached a concentration of 400 nM in the cytosol, corresponding to a 40% delivery efficiency; under equal conditions, ZF5.3-AGT54^Rho^ reached 139.2 nM, and ZF5.3-SNAP^Rho^ only reached 44.2 nM. It is notable that ZF5.3-GE-AGT^Rho^ and ZF5.3-DHFR^Rho^ show comparable thermal stabilities (*T_m_ values of 35°C and 33°C, respectively) but ZF5.3-DHFR^Rho^ reaches the cytosol significantly more efficiently under comparable incubation conditions; this likely relates to the relatively higher total uptake of ZF5.3-DHFR^Rho^ (Fig. 1 b,c). On their own, the series of SNAP variants lacking ZF5.3 reached the cytosol at virtually undetectable levels (cytosolic concentrations between 9 and 16.8 nM after a 30 min incubation) with minimal differences among the three (Fig. 3c,d), indicating that the relationship between thermostability and delivery is unique to a ZF5.3-driven pathway.

### ZF5.3-mediated delivery of a small but stable mini-protein

Membrane translocation machines that transit unfolded protein domains sometimes tolerate secondary structures or even folded proteins if they are small and compact (18, 38, 39). Moreover, proteins with high pI’s (excess cationic surface charge) can engage negatively charged phospholipids for enhanced cellular uptake (40, 41). Small stable protein domains, whether natural, evolved, or designed, are desirable research tools and are increasingly represented in clinical trials (42). Indeed, ZF5.3 was recently shown to facilitate cytosolic delivery of a nanobody-derived Bio-Protac that catalytically induces degradation of Bcl-11 and upregulates fetal hemoglobin production, although the delivery efficiency was not evaluated (9). To more quantitatively evaluate whether small, stable proteins could be delivered effectively by ZF5.3, we turned to synthetic mini-proteins derived from the fibronectin type III domain (monobodies). Monobodies can be engineered to display exceptionally high affinity for difficult-to-inhibit proteins (43, 44), are 20-25% more compact than nanobodies (44), and are not themselves cell permeant (45, 46). In particular, we focused on NS1 (Fig. 4a), a small (12 kD), cationic (pI = 9.2 when conjugated to ZF5.3) monobody that binds HRAS and KRAS with high affinity (K_D_ values of 15 nM and 65 nM, respectively), and inhibits KRAS-driven tumor growth when expressed in vivo (46).

**Fig. 4.**
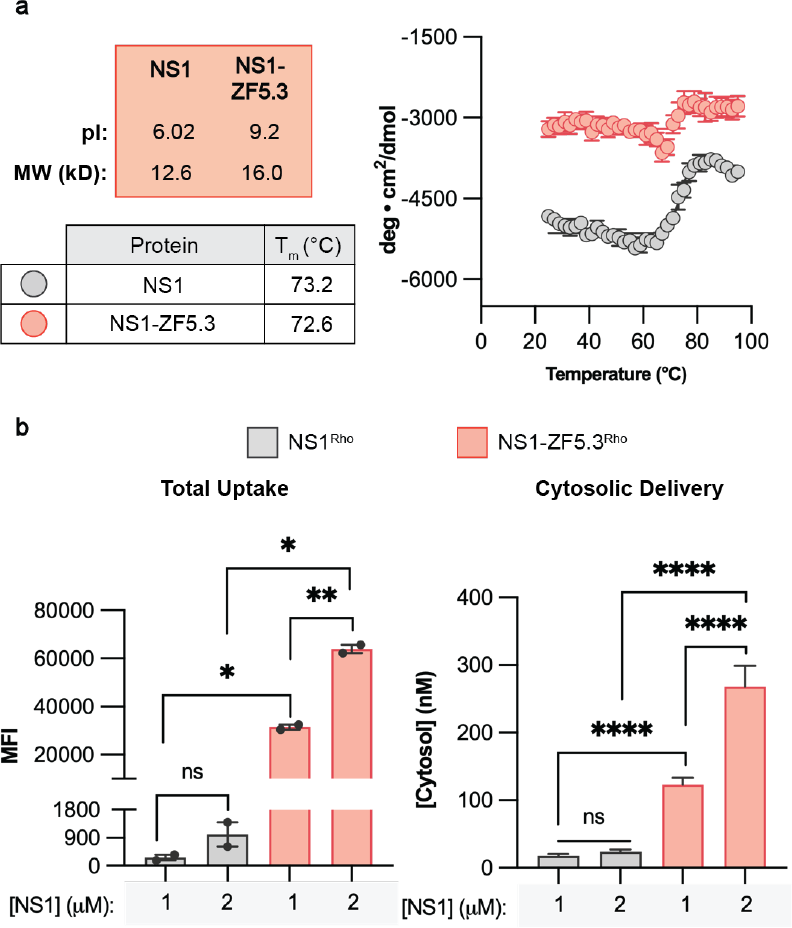
ZF5.3 can deliver the Ras-targeting monobody NS1 to the cytosol of cells. **a**, Predicted isoelectric point (pI), molecular weight (kD), and experimentally determined *T_m_ (°C) of the Ras-targeting monobody NS1, with or without ZF5.3. The temperature-dependent CD signal at 218 nm was measured in 2° increments between 25 and 90°C and the ellipticity was fitted using a Boltzmann sigmoidal nonlinear regression to obtain *T_m_ values. Each protein was measured in a buffer containing 20 mM Tris, 150 mM KCl, 0.5 mM TCEP pH 7.5. The observation that molar ellipticity decreases with temperature until ∼65°C for both NS1 and NS1-ZF5.3 has been documented for fibronectin-like domains (47, 48) and may be due to a partial loss in structure at low temperatures. **b**, Total cellular uptake (left) and cytosolic concentration (right) of NS1^Rho^ or NS1-ZF5.3^Rho^ were analyzed by flow cytometry and FCS. Saos-2 cells were incubated with 1 or 2 µM of the given protein for 1 h before the cellular workup and measurements as described previously. Flow cytometry values are provided as Median Fluorescence Intensity for the lissamine rhodamine B channel, n = 20000 total per condition with two biological replicates each (mean ± SEM). FCS values are provided in nM, n>25 for each FCS condition with two biological replicates each (mean ± SEM). Statistical significance was assessed using the Brown-Forsythe and Welch one-way analysis of variance (ANOVA) followed by an unpaired t-test with Welch’s correction. ****p ≤ 0.0001, ***p ≤ 0.001, **p ≤ 0.01, *p ≤ 0.05.

NS1 and NS1-ZF5.3 were expressed and purified, labeled at the C-terminus with rhodamine via a thiol-Michael addition reaction (*SI Appendix*, Fig. S8a,b), and characterized by LC/MS and CD (*SI Appendix*, Fig. S8c,d). Comparison of the wavelength spectra for NS1 and NS1-ZF5.3 suggests the addition of ZF5.3 does not significantly perturb the secondary structure of NS1. As expected, both NS1 and NS1-ZF5.3 are highly thermostable (Fig. 4a; *T_m_ = 73.2°C for NS1 and 72.6°C for NS1-ZF5.3). To evaluate delivery, Saos-2 cells were treated with 1 - 2 µM NS1^Rho^ and NS1-ZF5.3^Rho^ for 1 h, washed and trypsinized, and analyzed by flow cytometry and FCS (Fig. 4b). Both the total uptake of NS1-ZF5.3^Rho^ and its ability to reach the cytosol were substantially higher than that of NS1^Rho^ (Fig. 4b). The total uptake of NS1 was improved by 63 - 117-fold upon conjugation to ZF5.3, whereas delivery to the cytosol was improved by 7-12-fold. NS1-ZF5.3^Rho^ reached a maximal cytosolic concentration of 122.9 nM and 268.3 nM with a starting incubation concentration of 1 and 2 µM, respectively, yielding a delivery efficiency of 12.3-13.4%. Under equivalent conditions, this cytosolic concentration is roughly equal to that of the mid-stable SNAP variant ZF5.3-AGT54^Rho^ (Fig. 3d), which has a significantly lower *T_m_ (46°), but also a less cationic pI (8.8) and a larger molecular weight (23.6 kDa). It is notable that the total uptake for NS1-ZF5.3^Rho^ was 11.9-fold higher than ZF5.3-AGT54^Rho^, even though the cytosolic concentrations were nearly equal. The uptake of NS1-ZF5.3^Rho^ more closely resembles that of ZF5.3-DHFR^Rho^ under equivalent conditions (MFI = 31450 and 33209, respectively, Fig. 1b and Fig. 4b), but the cytosolic delivery of ZF5.3-DHFR^Rho^ was much more efficient (123 nM and 393 nM, respectively, Fig. 1c and Fig. 4b). These results suggest that, although a cationic surface charge and compact fold can result in modest cytosolic delivery, the specific step(s) at which ZF5.3 conjugates escape the endocytic pathway is most efficient for easily unfoldable proteins.

### HOPS provides a portal for efficient endosomal escape of easily unfolded proteins

Given the evidence that efficient ZF5.3-mediated membrane translocation demands protein unfolding, we next asked whether this delivery pathway makes use of endosomal machinery. We were specifically interested in the role of the HOPS and CORVET complexes, two essential hexameric tethering complexes involved in endosomal maturation events (49, 50). HOPS coordinates with SNARE proteins and a Rab GTPase to drive late endosome-lysosome fusion, while CORVET performs an analogous role for early endosomal fusion (Fig. 5a) (10, 51, 52). Previous work revealed that efficient endosomal escape of ZF5.3, both alone and when fused to the intrinsically disordered cargo MeCP2, requires HOPS but not CORVET, suggesting an escape portal is generated during or after endolysosomal fusion (5, 6). Whether this dependency extended to all ZF5.3 cargoes or only those that easily unfold remained unclear.

**Fig. 5.**
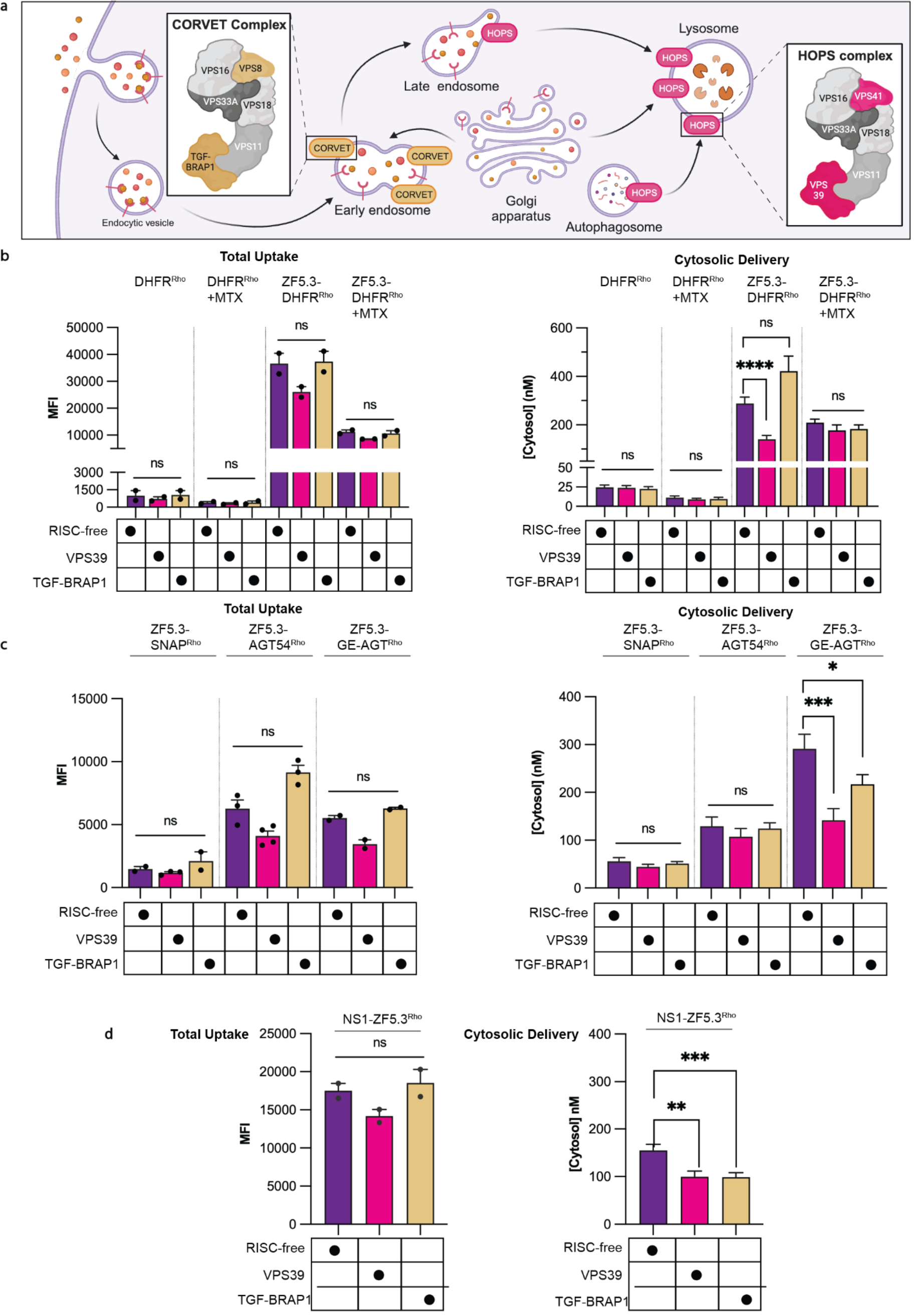
Easily unfolded proteins require HOPS and ZF5.3 to reach the cytosol efficiently. **a**, Knockdowns were performed for the VPS39 subunit of the HOPS complex or the TGF-BRAP1 subunit of the CORVET complex. Both complexes participate in membrane tethering for either Rab5+ early endosomes and maturing endosomes (CORVET) or Rab7+ and Lamp1+ late endosomes and lysosomes (HOPS). Schematic adapted from “Role of HOPS in Lysosome Formation”, by BioRender.com (2023). Retrieved from https://app.biorender.com/biorendertemplates. **b-d**, Plots illustrating the effects of VPS39 and TGF-BRAP1 knockdowns on total uptake (flow cytometry, Median Fluorescence Intensity) and cytosolic access (FCS, nM) for DHFR proteins (**b**), SNAP-tag variants (**c**), and NS1-ZF5.3 (**d**) relative to a RISC-free negative control. Two biological replicates were performed for each experiment, n = 20000 per condition in total for flow cytometry and n>15 per condition for FCS. Error bars represent the SEM. ****P < 0.0001, ***P < 0.001, **P < 0.01, *P < 0.05, not significant (ns) for P > 0.05 from one-way ANOVA with unpaired t-test with Welch’s correction.

We began by investigating the HOPS dependence of ZF5.3-mediated delivery of DHFR in the presence and absence of MTX. Saos-2 cells were transfected with siRNAs targeting either an essential HOPS subunit (VPS39) or the analogous CORVET subunit (TGF-BRAP1), as well as a non-targeting siRNA (RISC-free) as a negative control. All knockdowns were verified using qPCR (*SI Appendix*, Fig. S9). We then treated cells with 500 nM DHFR^Rho^ or ZF5.3-DHFR^Rho^ for 1 h and analyzed each sample by flow cytometry and FCS (Fig. 5b). Although depletion of VPS39 had only a modest effect on the total uptake of either DHFR^Rho^ or ZF5.3-DHFR^Rho^, it substantially (51%) decreased the efficiency with which ZF5.3-DHFR^Rho^ trafficked to the cytosol relative to the RISC-free control (Fig. 5b). Interestingly, knockdown of TGF-BRAP1 slightly increased the fraction of ZF5.3-DHFR^Rho^ that reached the cytosol (Fig. 5b), a pattern also observed for ZF5.3^Rho^ alone (5) but not for ZF5.3-MeCP2 (6). Notably, VPS39 knockdown had no effect on the cytosolic delivery of ZF5.3-DHFR^Rho^ in the presence of one equivalent MTX, nor any effect on delivery of DHFR^Rho^. These results demonstrate that ZF5.3-DHFR, like ZF5.3 alone and ZF5.3-MeCP2, makes use of late endosome tethering and/or fusion events to reach the cytosol. The lack of HOPS dependence for ZF5.3-DHFR^Rho^ in the presence of MTX, as well as DHFR^Rho^ (+/-MTX), suggests that certain proteins escape endosomes inefficiently through one or more pathways, but that attachment of ZF5.3 to a protein that easily unfolds biases endosomal escape toward a highly efficient, HOPS-dependent route.

To establish whether the link between HOPS and protein unfolding applied to other proteins, we examined the effect of HOPS- and CORVET-specific siRNA depletions on the uptake and cytosolic trafficking of SNAP-tag variants (Fig. 5c). As observed for DHFR^Rho^ and ZF5.3-DHFR^Rho^, depletion of VPS39 had no statistically significant effect on the uptake of any SNAP variant. Depletion of VPS39 also had no effect on the cytosolic delivery of the high-*Tm and mid-*Tm SNAP variants (ZF5.3-SNAP^Rho^ and ZF5.3-AGT54^Rho^) - in all cases the concentration established in the cytosol was relatively low (44 - 56 nM for ZF5.3-SNAP^Rho^ and 107 - 130 nM for ZF5.3- AGT54^Rho^). Depletion of VPS39 did, however, significantly decrease the cytosolic trafficking of the low-*Tm SNAP variant (ZF5.3-GE-AGT^Rho^), by 51.4%. Knockdown of TGF-BRAP1 had no effect on delivery of the high- and mid-*Tm variants and a mild but statistically significant decrease (25.5%) in delivery of the low-*Tm variant. The untagged SNAP^Rho^ variants reached extremely low cytosolic concentrations under all conditions tested and were too low to reliably quantify. For consistency, we also evaluated the effect of VPS39 and TGF-BRAP1 knockdown on NS1-ZF5.3^Rho^ delivery (Fig. 5c). Depletion of both VPS39 and TGF-BRAP1 had minimal effect on total uptake and a modest and statistically significant (36% and 37%, respectively) reduction in cytosolic concentration of NS1-ZF5.3^Rho^. The variable effect of TGF-BRAP1 knockdown on delivery of ZF5.3-DHFR^Rho^, ZF5.3-GE-AGT^Rho^, and NS1-ZF5.3^Rho^ likely indicates some complexity in how the endosomal maturation machinery is utilized. Together, these data suggest that ZF5.3 conjugates with easily unfolded cargos exploit a high-efficiency, HOPS-dependent pathway that can be partially adapted by cargos with high thermal stabilities provided the folded state is sufficiently compact and cationic. Even in this case, however, the delivery efficiency is markedly lower than that of a protein which can unfold under physiological conditions.

### STED microscopy reveals membrane-associated subcompartments within endolysosomes

But how does HOPS, which catalyzes homotypic and heterotypic membrane fusion from the cytosol, communicate with material within the endosomal lumen? Two lines of evidence suggest that endosomal escape involves more than the establishment of a membrane defect during vesicle fusion. First, efficient endosomal escape demands a covalent link between ZF5.3 and the delivered cargo (8). Second, ZF5.3 does not promote endosomal escape of other endosomally sequestered material (5). Although both ZF5.3 (5) and ZF5.3-DHFR localize primarily within the lumen of Lamp1+ endolysosomes when evaluated using confocal microscopy (*SI Appendix*, Fig. S10a), TauSTED microscopy of ZF5.3-DHFR^Rho^ treated cells (*SI Appendix*, Fig. S10b) revealed fluorescent populations that resemble intraluminal vesicles (ILVs, *SI Appendix*, Fig. S10c). ILVs are a critical component of multivesicular bodies, and it is possible that HOPS-catalyzed fusion events enable ZF5.3 and ZF5.3-DHFR to interact with ILVs in a manner that facilitates endosomal release. Notably, at super-resolution the fluorescent sub-populations all appear near endolysosomal membranes (*SI Appendix*, Fig. S10c), suggestive of membrane interactions that facilitate endosomal release along a concentration gradient into the cytosol. Precisely how luminal ZF5.3-DHFR communicates with HOPS to ultimately cross the endosomal membrane, and whether this mechanism definitively proceeds through an ILV-related pathway, is an area of active investigation.

## Discussion

Here we describe the first design rules for efficient endosomal escape of protein cargo. We find that the efficiency of ZF5.3-mediated protein delivery to the cytosol is highest when the protein cargo readily unfolds under physiological conditions. For cargos that are more stable, a compact size and cationic charge partially compensate to improve delivery efficiency, as observed for NS1-ZF5.3^Rho^. These results suggest that the impact of ZF5.3 as a delivery vehicle can be maximized by choosing cargos that adhere to these design rules (Fig. 6a,b). There are dozens of annotated proteins with T_m_ values comparable to those chosen in this study (53, 54), and hundreds of proteins containing >40% intrinsic disorder (55). Protein engineering efforts to introduce pH-or temperature-dependent destabilizing mutations into otherwise ideal therapeutic candidates to improve ZF5.3-mediated delivery, such as NS1, may also be a viable strategy to enhance delivery efficiency. The observation that ZF5.3 endosomal escape is most efficient when conjugated to low-*T_m_ proteins, and that this pathway demands communication between luminal ZF5.3 and cytosol-facing HOPS, suggests the existence of a selective portal through which membrane transport occurs. In nearly all cases, nature mediates such transport via a proteinaceous channel embedded within the membrane, such as the recently reported perforin-2 channel in dendritic cells (56). Whether ZF5.3 accomplishes its escape via lipid interactions or makes use of a yet-undetected protein channel remains under active investigation.

**Fig 6.**
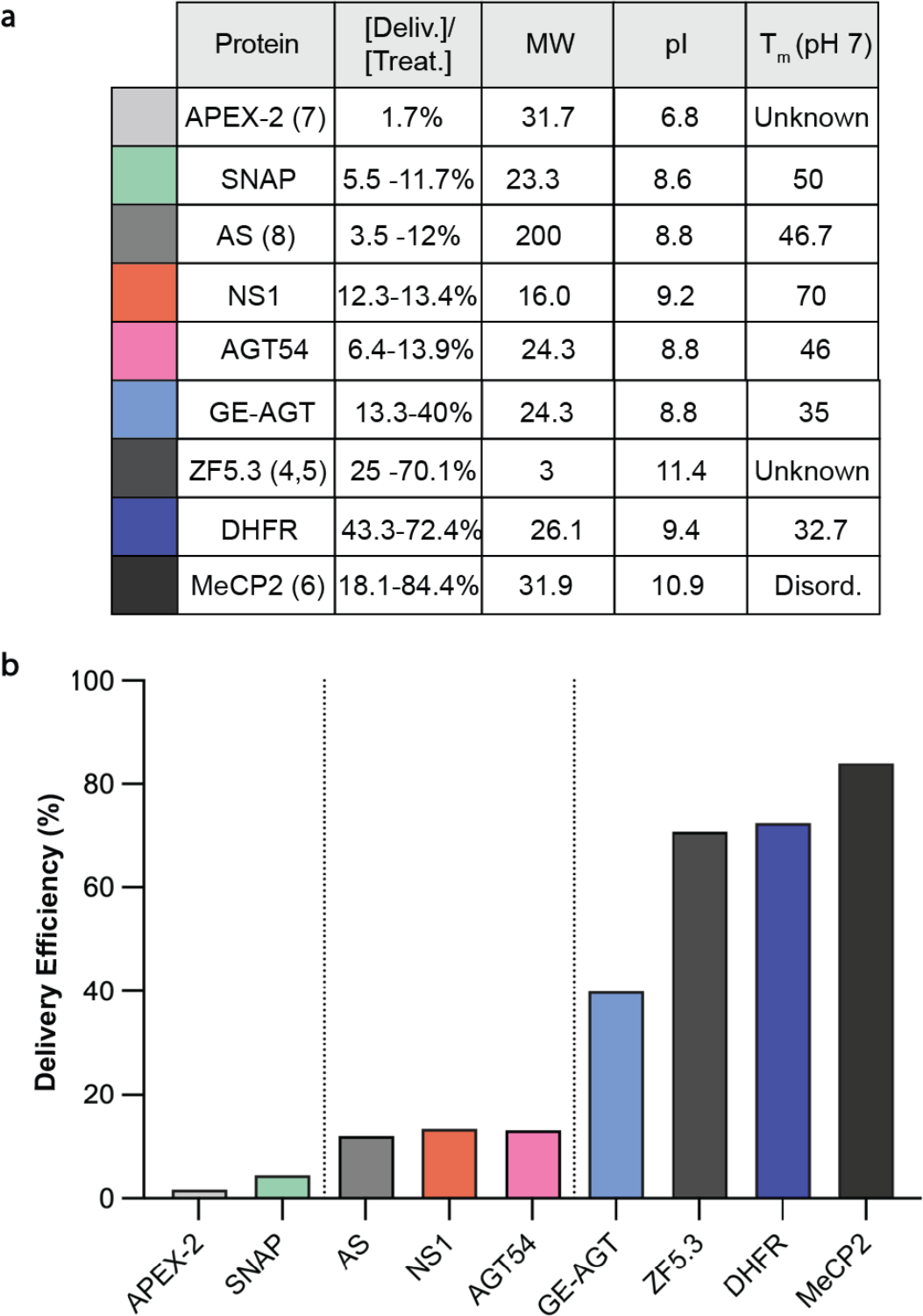
Efficient delivery of covalent ZF5.3 conjugates correlates directly with melting temperature. a, Biophysical parameters and delivery efficiency for ZF5.3 alone (dark gray) or when conjugated to various protein cargos. Delivery efficiency is defined as the concentration that reaches the cytosol or nucleus divided by the treatment concentration. Molecular weight (MW) is defined in kilodaltons, pI is the isoelectric point, and T_m_ is the melting temperature, determined experimentally when conjugated to ZF5.3. **b)** Graphical representation of the delivery efficiency for cargos listed in (a). Proteins with high T_m_ that are larger than 20 kD are delivered with the lowest efficiency. Proteins with a T_m_ ∼46, or a high T_m_ but small molecular weight, are delivered with mid-range efficiency. Only proteins with a T_m_ <35°C or that are intrinsically disordered are delivered with the highest efficiency.

## Materials & Methods

A detailed description of all materials and methods is provided in the *SI Appendix*.

## Acknowledgments

This work was supported by the National Science Foundation (Grant No. 2203903), the NIH (Grant No. R35GM134963), and Merck. L.L. was supported in part by a National Science Foundation Graduate Research Fellowship (Grant No. 2146752). A.S. is a Chan-Zuckerberg Biohub-San Francisco Investigator.

## References

1. B. Leader, Q. J. Baca, D. E. Golan, Protein therapeutics: a summary and pharmacological classification. Nat. Rev. Drug Discov. 7, 21–39 (2008).

2. C. Hald Albertsen, et al., The role of lipid components in lipid nanoparticles for vaccines and gene therapy. Adv. Drug Deliv. Rev. 188, 114416 (2022).

3. J. S. Appelbaum, et al., Arginine Topology Controls Escape of Minimally Cationic Proteins from Early Endosomes to the Cytoplasm. Chem. Biol. 19, 819–830 (2012).

4. J. R. LaRochelle, G. B. Cobb, A. Steinauer, E. Rhoades, A. Schepartz, Fluorescence Correlation Spectroscopy Reveals Highly Efficient Cytosolic Delivery of Certain Penta-Arg Proteins and Stapled Peptides. J. Am. Chem. Soc. 137, 2536–2541 (2015).

5. A. Steinauer, et al., HOPS-dependent endosomal fusion required for efficient cytosolic delivery of therapeutic peptides and small proteins. Proc. Natl. Acad. Sci. 116, 512–521 (2019).

6. X. Zhang, et al., Dose-Dependent Nuclear Delivery and Transcriptional Repression with a Cell-Penetrant MeCP2. ACS Cent. Sci. 9, 277–288 (2023).

7. R. F. Wissner, A. Steinauer, S. L. Knox, A. D. Thompson, A. Schepartz, Fluorescence Correlation Spectroscopy Reveals Efficient Cytosolic Delivery of Protein Cargo by Cell-Permeant Miniature Proteins. ACS Cent. Sci. 4, 1379–1393 (2018).

8. S. L. Knox, R. Wissner, S. Piszkiewicz, A. Schepartz, Cytosolic Delivery of Argininosuccinate Synthetase Using a Cell-Permeant Miniature Protein. ACS Cent. Sci. 7, 641–649 (2021).

9. F. Shen, et al., A Cell-Permeant Nanobody-Based Degrader That Induces Fetal Hemoglobin. ACS Cent. Sci. 8, 1695–1703 (2022).

10. H. Song, A. S. Orr, M. Lee, M. E. Harner, W. T. Wickner, HOPS recognizes each SNARE, assembling ternary trans-complexes for rapid fusion upon engagement with the 4th SNARE. eLife 9, e53559 (2020).

11. R. A. Arkowitz, J. C. Joly, W. Wickner, Translocation can drive the unfolding of a preprotein domain. EMBO J. 12, 243–253 (1993).

12. J. A. Lycklama a Nijeholt, A. J. M. Driessen, The bacterial Sec-translocase: structure and mechanism. Philos. Trans. R. Soc. B Biol. Sci. 367, 1016–1028 (2012).

13. M. Eilers, G. Schatz, Binding of a specific ligand inhibits import of a purified precursor protein into mitochondria. Nature 322, 228–232 (1986).

14. U. Wienhues, et al., Protein folding causes an arrest of preprotein translocation into mitochondria in vivo. J. Cell Biol. 115, 1601–1609 (1991).

15. D. Vestweber, J. Brunner, A. Baker, G. Schatz, A 42K outer-membrane protein is a component of the yeast mitochondrial protein import site. Nature 341, 205–209 (1989).

16. P. K. Kim, E. H. Hettema, Multiple Pathways for Protein Transport to Peroxisomes. J. Mol. Biol. 427, 1176–1190 (2015).

17. C. Rabouille, V. Malhotra, W. Nickel, Diversity in unconventional protein secretion. J. Cell Sci. 125, 5251–5255 (2012).

18. D. Pei, R. E. Dalbey, Membrane translocation of folded proteins. J. Biol. Chem. 298, 102107 (2022).

19. C. Chávez-García, J. Hénin, M. Karttunen, Multiscale Computational Study of the Conformation of the Full-Length Intrinsically Disordered Protein MeCP2. J. Chem. Inf. Model. 62, 958–970 (2022).

20. N. Salvador, C. Aguado, M. Horst, E. Knecht, Import of a Cytosolic Protein into Lysosomes by Chaperone-mediated Autophagy Depends on Its Folding State*. J. Biol. Chem. 275, 27447–27456 (2000).

21. F. A. Agarraberes, J. F. Dice, A molecular chaperone complex at the lysosomal membrane is required for protein translocation. J. Cell Sci. 114, 2491–2499 (2001).

22. J. Shi, et al., A technique for delineating the unfolding requirements for substrate entry into retrotranslocons during endoplasmic reticulum–associated degradation. J. Biol. Chem. 294, 20084–20096 (2019).

23. B. Beaumelle, M.-P. Taupiac, J. M. Lord, L. M. Roberts, Ricin A Chain Can Transport Unfolded Dihydrofolate Reductase into the Cytosol*. J. Biol. Chem. 272, 22097–22102 (1997).

24. O. Klingenberg, S. Olsnes, Ability of methotrexate to inhibit translocation to the cytosol of dihydrofolate reductase fused to diphtheria toxin. Biochem. J. 313, 647–653 (1996).

25. G. Haug, et al., Cellular uptake of Clostridium botulinum C2 toxin: membrane translocation of a fusion toxin requires unfolding of its dihydrofolate reductase domain. Biochemistry 42, 15284–15291 (2003).

26. S. A. Kim, K. G. Heinze, P. Schwille, Fluorescence correlation spectroscopy in living cells. Nat. Methods 4, 963–973 (2007).

27. S. L. Knox, et al., “Chapter Twenty-One - Quantification of protein delivery in live cells using fluorescence correlation spectroscopy” in Methods in Enzymology, Chemical Tools for Imaging, Manipulating, and Tracking Biological Systems: Diverse Chemical, Optical and Bioorthogonal Methods., D. M. Chenoweth, Ed. (Academic Press, 2020), pp. 477–505.

28. D. Magde, E. Elson, W. W. Webb, Thermodynamic Fluctuations in a Reacting System— Measurement by Fluorescence Correlation Spectroscopy. Phys. Rev. Lett. 29, 705–708 (1972).

29. P. Schwille, Fluorescence correlation spectroscopy and its potential for intracellular applications. Cell Biochem. Biophys. 34, 383–408 (2001).

30. E. L. Elson, Fluorescence Correlation Spectroscopy: Past, Present, Future. Biophys. J. 101, 2855–2870 (2011).

31. M. C. Waltham, J. W. Holland, P. F. Nixon, D. J. Winzor, Thermodynamic characterization of the interactions of methotrexate with dihydrofolate reductase by quantitative affinity chromatography. Biochem. Pharmacol. 37, 541–545 (1988).

32. W. C. Werkheiser, Specific Binding of 4-Amino Folic Acid Analogues by Folic Acid Reductase. J. Biol. Chem. 236, 888–893 (1961).

33. R. S. Swanwick, A. M. Daines, L.-H. Tey, S. L. Flitsch, R. K. Allemann, Increased Thermal Stability of Site-Selectively Glycosylated Dihydrofolate Reductase. ChemBioChem 6, 1338–1340 (2005).

34. B. Mollwitz, et al., Directed Evolution of the Suicide Protein O6-Alkylguanine-DNA Alkyltransferase for Increased Reactivity Results in an Alkylated Protein with Exceptional Stability. Biochemistry 51, 986–994 (2012).

35. A. Juillerat, et al., Directed Evolution of O6-Alkylguanine-DNA Alkyltransferase for Efficient Labeling of Fusion Proteins with Small Molecules In Vivo. Chem. Biol. 10, 313–317 (2003).

36. A. Juillerat, et al., Engineering Substrate Specificity of O6-Alkylguanine-DNA Alkyltransferase for Specific Protein Labeling in Living Cells. ChemBioChem 6, 1263–1269 (2005).

37. T. Gronemeyer, C. Chidley, A. Juillerat, C. Heinis, K. Johnsson, Directed evolution of O6-alkylguanine-DNA alkyltransferase for applications in protein labeling. Protein Eng. Des. Sel. 19, 309–316 (2006).

38. E. A. Craig, Hsp70 at the membrane: driving protein translocation. BMC Biol. 16, 11 (2018).

39. I. H. Madshus, S. Olsnes, H. Stenmark, Membrane translocation of diphtheria toxin carrying passenger protein domains. Infect. Immun. 60, 3296–3302 (1992).

40. D. B. Thompson, R. Villaseñor, B. M. Dorr, M. Zerial, D. R. Liu, Cellular Uptake Mechanisms and Endosomal Trafficking of Supercharged Proteins. Chem. Biol. 19, 831–843 (2012).

41. F. Madani, S. Lindberg, Ü. Langel, S. Futaki, A. Gräslund, Mechanisms of Cellular Uptake of Cell-Penetrating Peptides. J. Biophys. 2011, 414729 (2011).

42. B. Jin, S. Odongo, M. Radwanska, S. Magez, Nanobodies: A Review of Generation, Diagnostics and Therapeutics. Int. J. Mol. Sci. 24, 5994 (2023).

43. A. Koide, C. W. Bailey, X. Huang, S. Koide, The fibronectin type III domain as a scaffold for novel binding proteins11Edited by J. Wells. J. Mol. Biol. 284, 1141–1151 (1998).

44. C. Carrasco-López, et al., Development of light-responsive protein binding in the monobody non-immunoglobulin scaffold. Nat. Commun. 11, 4045 (2020).

45. F. Sha, G. Salzman, A. Gupta, S. Koide, Monobodies and other synthetic binding proteins for expanding protein science. Protein Sci. 26, 910–924 (2017).

46. R. Spencer-Smith, et al., Inhibition of RAS function through targeting an allosteric regulatory site. Nat. Chem. Biol. 13, 62–68 (2017).

47. S. V. Litvinovich, et al., Formation of amyloid-like fibrils by self-association of a partially unfolded fibronectin type III module11Edited by R. Huber. J. Mol. Biol. 280, 245–258 (1998).

48. M. A. Kruziki, S. Bhatnagar, D. R. Woldring, V. T. Duong, B. J. Hackel, A 45-Amino-Acid Scaffold Mined from the PDB for High-Affinity Ligand Engineering. Chem. Biol. 22, 946–956 (2015).

49. S. Messler, et al., The TGF-β signaling modulators TRAP1/TGFBRAP1 and VPS39/Vam6/TLP are essential for early embryonic development. Immunobiology 216, 343–350 (2011).

50. H. Luo, et al., DEG 15, an update of the Database of Essential Genes that includes built-in analysis tools. Nucleic Acids Res. 49, D677–D686 (2021).

51. R. van der Kant, et al., Characterization of the Mammalian CORVET and HOPS Complexes and Their Modular Restructuring for Endosome Specificity*. J. Biol. Chem. 290, 30280–30290 (2015).

52. D. Shvarev, et al., Structure of the HOPS tethering complex, a lysosomal membrane fusion machinery. eLife 11, e80901 (2022).

53. R. Nikam, A. Kulandaisamy, K. Harini, D. Sharma, M. M. Gromiha, ProThermDB: thermodynamic database for proteins and mutants revisited after 15 years. Nucleic Acids Res. 49, D420–D424 (2021).

54. A. Jarzab, et al., Meltome atlas—thermal proteome stability across the tree of life. Nat Methods 17, 495–503 (2020).

55. F. Quaglia, et al., DisProt in 2022: improved quality and accessibility of protein intrinsic disorder annotation. Nucleic Acids Res. 50, D480–D487 (2022).

56. P. Rodríguez-Silvestre, et al., Perforin-2 is a pore-forming effector of endocytic escape in cross-presenting dendritic cells. Science 380, 1258–1265 (2023).

