## Supplementary material for "Design rules for efficient endosomal escape": SI Appendix

### SI Appendix for

* Alanna Schepartz

**Author Contributions:** Study conception and design: M.Z. and A.S.; preparation of materials: M.Z., A.V., X.Z. and L.L.; data collection: M.Z., A.V., X.Z., and N.D.; analysis and interpretation of results: M.Z., A.V., and N.D.; and manuscript preparation: M.Z. and A.S.

**Competing Interest Statement:** None.

**Classification:** Major: Biological Sciences; Minor: Biochemistry

**Keywords:** protein delivery, protein folding, protein therapeutics, fluorescence correlation spectroscopy, nanobody.

**This PDF file includes:**

Reagents & Chemicals
Detailed Methods
Supplementary Figures 1-10
Supplementary Tables 1-3
SI References

**Reagents & Chemicals:**

Cell culture/Microscopy: SsoFast$^{TM}$ EvaGreen® Supermix (#1725200, BioRad); McCoy's 5A Medium without phenol red (#SH30270, Cytiva); all siRNAs (Dharmacon, **Table S1**); RT-qPCR primers (IDT, **Table S1**); 4-well #1.5H glass bottom coverslip (#80427, Ibidi); Lipofectamine RNAiMAX transfection reagent (#13778150, Invitrogen); Fibronectin bovine plasma (#F1141), Omni 0.5mL Tubes w/ 1.4mm Ceramic Beads (#19626) (Millipore Sigma); FuGENE HD transfection reagent (Promega, #E2311); Gibco$^{TM}$ Dulbecco's modified eagle medium (DMEM) without phenol red, high glucose, with 25 mM HEPES (#21063), Gibco$^{TM}$ GlutaMAX$^{TM}$ Supplement, 200 mM (#35050061), Gibco$^{TM}$ Sodium Pyruvate, 100 mM (#11360070), Gibco$^{TM}$ TrypLE Express Enzyme (1x) with (#12605010) and without (#12604013) phenol red, McCoy's 5a Medium (#16600082), Nunc™ Lab-Tek™ 8-well Chambered Coverglass (#155411) (Thermofisher Scientific); Saos-2 cell stock (UC Berkeley Cell Culture Facility)

Cell lines & plasmids: CMV-LAMP1-HaloTag plasmid (#164209, Addgene); *E. coli* BL21(DE3) competent cells (#230132) and *E. coli* BL21-CodonPlus(DE3)-RP cells (#230255) (Agilent); all synthetic gBlocks (IDT, **Table S1**); pet-32a(+) vector (Millipore Sigma, #69015)

Chemicals: O6-(4-Aminomethyl-benzyl)guanine (#12560, AAT Bioquest); Isopropyl β-D-thiogalactopyranoside (#367-93-1, American Bio); Ammonium Chloride (#A9434), Biotin (#B4501), Calcium Chloride (#C5670), Fmoc-Gly-OH (#47627), Fmoc-Lys(Mtt)-OH (#852065), Glucose (#G5767), Lissamine Rhodamine B Sulfonyl Chloride (#86186), M9 Minimal Salts 5x (#M6030), Magnesium Sulfate (#M2643), N,N-Diisopropylethylamine (#387649), Potassium Chloride (#P3911), Sodium Chloride (#S9888) (Millipore Sigma); Lissamine Rhodamine B C2 Maleimide (Tenova, #T01196); 1M Tris-HCl buffer pH 7.5 (#15567027), Ammonium Sulfate (#011566), Glycerol (#J61059), Imidazole (#A10221), Invitrogen Alexa Fluor$^{TM}$ 594 hydrazide (#A10438), Rhodamine Red™ C2 Maleimide (#R6029), Thiamine Hydrochloride (#148990100), Tris (2-Carboxyethyl) Phosphine Hydrochloride (#50-153-2844) (Thermofisher Scientific)

Molecular biology reagents: α-Tubulin Rabbit antibody (#2125S), Anti-rabbit IgG, HRP-linked Antibody (#7074S), His-tag Rabbit antibody (#2365S), Lamp1 XP® Rabbit antibody (#9091S) (Cell Signaling Technology); HiTrap® SP HP 5-mL column (#17115101), StrepTrap® HP 5-mL column (#28907547), PD-10 desalting columns (#17085101) (Cytiva); 3 kDa MWCO Amicon® Ultra-15 Centrifugal Filter Unit (#UFC90003), 10 kDa MWCO Amicon® Ultra-15 Centrifugal Filter Unit (#UFC9010), Benzonase® Nuclease HC (#71205), cOmplete, Mini EDTA-free protease inhibitor cocktail (#4693159001), ZipTips with 0.6 µL C4 resin (#ZTC04S008), 0.22 µm hydrophilic PVDF membrane filter (#SLGVR33RS, Millipore Sigma); NEBuilder® HiFi DNA Assembly Master Mix (#E2621L), Q5® High-Fidelity 2X Master Mix (#M0492S) (New England Biolabs); TALON® metal affinity resin (Takara, #635504); Pierce™ 660 nm Protein Assay Reagent (#22660), Pierce™ BCA Protein Assay Kit (#23227), Pierce™ Universal Nuclease for Cell Lysis (#88700) (Thermofisher Scientific)

#### Detailed Methods

#### Plasmid construction

All gBlocks were codon-optimized for expression in E. coli B strain and ordered from Integrated DNA Technologies (IDT). The gBlocks for DHFR proteins encoded the DNA sequence for murine DHFR with or without an N-terminal ZF5.3 separated from the DHFR sequence with a glycine-serine-glycine (GSG) linker. Both sequences included a C-terminal LPETGG motif for sortase conjugation (1–3) followed by a His6 tag for affinity purification. gBlocks encoding AGT54 and GE-AGT (4) were designed with or without ZF5.3 at the N-terminus separated by a GSG linker and included a C-terminal His6 tag. The plasmids for SNAP-tag and ZF5.3-SNAP-tag were reported in a previous publication (1). For NS1 (5), initial attempts to purify a variant with ZF5.3 at the N-terminus were unsuccessful and could not be optimized. gBlocks encoding the sequence for NS1 with or without a C-terminal ZF5.3 separated by a $(GGGGS)_3$ linker were designed with an N-terminal Strep tag for affinity purification and a C-terminal cysteine residue for fluorophore conjugation. All gBlocks were inserted into a pET-32a(+) backbone vector (Sigma) linearized with **Primers 1** and **2** using Gibson assembly. The identities of all plasmids were confirmed by Sanger and whole plasmid sequencing. Relevant DNA and protein sequences are listed in **Table S1**.

#### Protein expression and purification

The expression and purification protocols for all proteins used in this study are detailed below. For all expression steps, the cultures were grown in an incubator at the specified temperature with shaking at 200 rpm. The compositions of all buffers described in this section are listed in **Table S2**.

##### Expression and purification of DHFR proteins

The plasmids encoding DHFR and ZF5.3-DHFR were transformed into E. coli BL21(DE3) competent cells and selected on an ampicillin (amp) LB agar plate. For each protein, one colony was used to inoculate a 20 mL overnight starter culture of LB media supplemented with 100 µg/mL amp and incubated at 37°C. The next morning, the starter culture was added to 1 L of LB with 100 mg/L amp and incubated at 37°C until the optical density ($OD_{600}$) reached 0.6-0.8. Protein expression was induced with 1 mM IPTG and the cultures were transferred to 20°C for 20 h. All steps from this point forward were performed at 4°C. To harvest the cells, cultures were spun at 4300 g for 40 min and resuspended in 20 mL of ice-cold Lysis Buffer 1 supplemented with 1 cOmplete, mini EDTA-free protease inhibitor cocktail tablet and 1 µL of Pierce Universal Nuclease for Cell Lysis. Cells were lysed by sonication for 7 min total, pulsing for 30 s and recovering for 30 s for 7 cycles. The lysate was cleared by centrifugation at 18,000 rpm for 35 min and transferred to a 50 mL conical tube containing 2 mL of TALON® metal affinity resin pre-equilibrated with Wash Buffer 2. The lysate was incubated with the resin for 40 min with gentle rotation. After incubation, the mixture was transferred to a gravity column and the flowthrough was allowed to drain. The resin was washed twice with 20 mL of Wash Buffer 1 and twice with 20 mL of Wash Buffer 2. The protein was then eluted with 14 mL of Elution Buffer 1 and collected in 1 mL fractions. All fractions were run on an SDS-PAGE gel, and fractions containing pure protein were pooled and dialyzed overnight at 4°C into 1 L of Storage Buffer 1, supplemented with 100 µM $ZnCl_2$ for ZF5.3-DHFR. The next day, the protein was quantified using a Pierce™ BCA Protein Assay kit and, if needed, concentrated to >1 mg/mL using an Amicon 10 kDA MWCO spin concentrator. The identities of final purified proteins were confirmed using liquid chromatography-mass spectrometry (Agilent Infinity II LC/6530 Accurate- Mass Q-TOF LC/MS) and stored at -80° until use.

##### Expression and purification of sortase enzymes

Two variants of the hepta-mutant Staphylococcus aureus Sortase A (SrtA7m) (6) were used for conjugation of a rhodamine dye to the C-terminus of DHFR proteins: one encoding S35 StrepTagII-SrtA7m-His6 (for generation of $DHFR^{Rho}$), and one encoding His6-SUMO-SrtA7m (for generation of ZF5.3-DHFR$^{Rho}$). Purification of both sortase enzymes was performed as previously described (3) with minor modifications. Both plasmids were transformed into E. coli BL21(DE3) competent cells and selected on a 50 μg/mL kanamycin (kan) LB agar plate (for S35 StrepTagII-SrtA7m-His6) or a 100 μg/mL amp LB agar plate (His6-SUMO-SrtA7m). A 5 mL starter culture inoculated with 1 colony was prepared for each with the appropriate antibiotic and grown at 37°C for 3.5 h. The starter culture was diluted into 1 L of LB with antibiotic and grown to $OD_{600}$ = 0.6 - 0.8, followed by induction with 0.5 mM IPTG. The culture was then transferred to 30°C for 18-24 h. All steps from this point forward were performed at 4°C. Cells were harvested by centrifugation at 4300 g for 40 min, resuspended in 20 mL Lysis Buffer 1, and lysed by sonication for 3 min (30 s on/30 s off pulses). The lysate was cleared by centrifugation at 10,000 rpm for 40 min and transferred to a 50 mL Falcon tube containing 2 mL of TALON® metal affinity resin pre-equilibrated with Wash Buffer 1. The resin was incubated with cleared lysate for 1 h with gentle rotation and added to a gravity column. The flowthrough was collected and the column was washed 3 times with 20 mL of Wash Buffer 1. The protein was then eluted in 13 mL of Elution Buffer 1, collected in 1 mL fractions, and run on an SDS-PAGE gel. Fractions containing pure protein were pooled and dialyzed overnight into Storage Buffer 1. The next day, the protein was quantified using a Pierce™ BCA Protein Assay kit.

For His6-SUMO-SrtA7m, the His6-SUMO was then cleaved by incubation with His6-SUMO protease (purified as previously described (2)) at a 4:1 molar ratio of His6-SUMO-SrtA7m to SUMO protease at room temperature for 2 hr. The cleaved SrtA7m protein was purified from the reaction mixture by incubation with TALON® metal affinity resin for 1 h at 4°C followed by collection of the column flowthrough and 10 1-mL wash fractions. The fractions were analyzed by SDS-PAGE, and those containing cleaved SrtA7m protein were pooled and dialyzed into fresh Storage Buffer 1 overnight at 4°C. Protein concentration was determined using a Pierce™ BCA Protein Assay kit. If needed, proteins were concentrated to >3 mg/mL with an Amicon 3 kDa MWCO spin concentrator and stored at -80°C until use.

Expression and purification of SNAP-tag proteins

*Expression/Purification of SNAP-tag, AGT54, and GE-AGT*:
The plasmids encoding SNAP-tag, AGT54, and GE-AGT were transformed into E. coli BL21(DE3) competent cells and selected on an amp/LB agar plate. For each protein, one colony was used to inoculate a 20 mL overnight starter culture of LB media supplemented with 100 μg/mL amp grown at 37°C. The next morning, the starter cultures were added to 1 L of LB per protein with 100 mg/L amp at 37°C until $OD_{600}$ = 0.6 - 0.8. Protein expression was induced with 1 mM IPTG, and the cultures were transferred to 18°C for 18 - 22 h. All steps from this point forward were performed at 4°C. The cells were harvested by centrifugation at 4300 g for 40 min and pellets were resuspended in 20 mL Lysis Buffer 2 supplemented with 1 cOmplete, mini EDTA-free protease inhibitor cocktail tablet per pellet. The cells were lysed using an EmulsiFlex-C3 homogenizer, and the lysate was cleared by centrifugation at 18,000 rpm for 35 min at 4°C. The cleared lysate was incubated with 2 mL of TALON® metal affinity resin for 1 h at 4°C with gentle rotation. The mixture was then added to a gravity column and the flowthrough was collected. The column was washed twice with High-Salt Buffer and twice with Low-Salt Buffer. The protein was then eluted with 10 mL Elution Buffer 2 and analyzed by SDS-PAGE. Fractions containing pure protein were pooled and dialyzed into Storage Buffer 2 overnight at 4°C. We observed that protein concentration using standard Amicon spin concentrators resulted in

significant sample loss, so the proteins were concentrated using a 65% w/v ammonium sulfate precipitation (7). Ammonium sulfate was added to the protein solution in a 15 mL conical tube, vortexed until fully dissolved, and incubated with rotation at 4°C for 10 min. The solution was then centrifuged at 4300 g for 10 min to sediment the precipitated protein in the pellet. The protein was resuspended in Storage Buffer 2. Protein concentration was determined using the Beer-Lambert law $A = \varepsilon bC$ where A = absorbance at 280 nm measured on a NanoDrop spectrophotometer (ND-1000), ε = the extinction coefficient for the protein, b = the optical path length in cm, and C = the protein concentration in M. To confirm the identity of all proteins, ~5 μg of each was desalted using ZipTip$_{C4}$® pipette tips (EMD Millipore) and analyzed by LC/MS. Purified proteins were stored at -80°C until use.

*Expression of ZF5.3-SNAP-tag*:
The expression of ZF5.3-SNAP was modified slightly from previous protocols to enable expression in minimal media in order to prevent formation of a +80 Da oxidative modification (1). The plasmid encoding ZF5.3-SNAP was transformed into E. coli BL21(DE3) competent cells and selected on an amp/LB agar plate. One colony was used to inoculate a 5 mL starter culture in LB supplemented with 100 μg/mL amp, which was grown for 6 h at 37°C. The starter culture was then transferred to 500 mL minimal M9 media supplemented with 1 g/L $NH_4Cl$, 4 g/L glucose, 2 mM $MgSO_4$, 0.1 mM $CaCl_2$, 100 mg/L amp, and 1 mM thiamine and grown with shaking overnight at 37°C. The next morning, 50 mL of overnight culture was transferred to 1 L of minimal M9 media with the same supplements described above plus a dash of biotin. The culture was grown with shaking at 37°C until $OD_{600}$ = 0.6, followed by induction with 1 mM IPTG. The culture was then transferred to 18°C and grown for 18 - 22 h.

*Expression of ZF5.3-AGT54*:
The expression of ZF5.3-AGT54 followed the same protocol as the untagged variants. The plasmid encoding ZF5.3-AGT54 was transformed into E. coli BL21(DE3) competent cells and selected on an amp/LB agar plate. One colony was used to inoculate a 20-mL overnight starter culture of LB media supplemented with 100 μg/mL amp. The next morning, the starter culture was added to 1 L of LB with 100 mg/L amp and incubated at 37°C until $OD_{600}$ = 0.6 - 0.8. Protein expression was induced with 1 mM IPTG, and the cultures were transferred to 18°C for 18 - 22 h.

*Expression of ZF5.3-GE-AGT*:
The expression of ZF5.3-GE-AGT was optimized further to improve yield. The plasmid encoding ZF5.3-GE-AGT was transformed into E. coli BL21-CodonPlus(DE3)-RP cells and selected on a double antibiotic LB agar plate containing 100 μg/mL amp and 25 μg/mL chloramphenicol (Cm). One colony was used to inoculate a 20 mL overnight starter culture of LB media supplemented with 100 μg/mL amp and 25 g/mL Cm, which was grown at 37°C. The starter culture was added to 1 L of LB with the appropriate antibiotics and grown to $OD_{600}$ = 0.8. ZF5.3-GE-AGT expression was induced with 0.5 mM IPTG and transferred to 18°C for 6 h.

*Purification of ZF5.3-SNAP, ZF5.3-AGT54, and ZF5.3-GE-AGT*:
After expression, all three proteins were purified using the same protocol. All steps were carried out at 4°C. Cells were harvested by centrifugation at 4300 g for 40 min, and pellets were resuspended in 20 mL Lysis Buffer 2 supplemented with 1 cOmplete, mini EDTA-free protease inhibitor cocktail tablet per pellet. The cells were lysed using an EmulsiFlex-C3 homogenizer, and the lysate was cleared by centrifugation at 18,000 rpm for 35 min at 4°C. The cleared lysate was

passed through a 0.22 μm membrane filter and the protein of interest was isolated using a two-step purification. First, the filtered lysate was applied to a 5 mL HiTrap® SP HP cation exchange column (Cytiva) and eluted using a linear gradient from 0.15 - 1 M NaCl over 10 column volumes (CV). Protein-containing fractions were analyzed by SDS-PAGE, and those containing the desired protein were pooled and further purified using TALON® metal affinity resin as described for SNAP-tag, AGT54, and GE-AGT above. We observed that the ZF5.3-tagged variants were not stable for overnight dialysis, and instead they were concentrated using a 65% w/v ammonium sulfate precipitation directly after elution from the resin. The precipitated protein was resuspended in Storage Buffer 2 and quantified using absorbance at 280 nm as described above. To confirm the identity of all proteins, ~5 μg of each was desalted using ZipTip$_{C4}$® pipette tips and analyzed by LC/MS. Purified proteins were stored at -80°C until use.

Expression and purification of NS1 proteins

*Expression/purification of NS1 and NS1-ZF5.3:*
The plasmid encoding NS1 or NS1-ZF5.3 was transformed into E. coli BL21(DE3) competent cells and selected on an amp/LB agar plate. One colony was used to inoculate a 5 mL starter culture of LB media supplemented with 100 μg/mL amp, which was grown at 37°C overnight. The next day, the starter culture was added to 1 L of LB with 100 mg/L amp and incubated with shaking at 37°C until the $OD_{600}$ reached 0.6-0.7. Expression was induced with 0.5 mM IPTG and, for NS1-ZF5.3 only, $ZnCl_2$ was added to the LB at a final concentration of 100 μM. The cultures were transferred to 18°C for 14-16 h.

All steps from this point forward were performed at 4°C. The cells were harvested by centrifugation at 4300 g for 30 min and resuspended in 10 mL ice-cold Lysis Buffer 3 supplemented with 1 μL of Benzonase® Nuclease and 1 cOmplete, mini EDTA-free protease inhibitor cocktail tablet. Cells were lysed by sonicating for 30 s and recovering for 30 s for 7 cycles. The lysate was cleared by centrifugation at 16,000 rpm for 35 min, filtered with a 0.22 μm membrane, and subjected to a one-step (for NS1) or two-step (for NS1-ZF5.3) purification. For NS1, the filtered lysate was applied to a 5 mL StrepTrap® HP Streptactin Sepharose affinity column (Cytiva) and eluted with a linear gradient from 0 - 2.5 mM desthiobiotin over 10 CV. Fractions containing pure protein were assessed using SDS-PAGE. For NS1-ZF5.3, the filtered lysate was first applied to a 5 mL HiTrap® SP HP cation exchange column (Cytiva) and eluted using a linear gradient from 0.05 - 2 M NaCl over 10 CV. Protein-containing fractions were analyzed by SDS-PAGE. We observed significant nucleic acid contamination for some fractions (monitored using a NanoDrop spectrophotometer) and only chose fractions with a 260/280 nm ratio < 0.6 for the next purification step. These fractions were then applied to a 5 mL StrepTrap® HP Streptactin Sepharose affinity column (Cytiva) and eluted with a linear gradient from 0 - 2.5 mM desthiobiotin over 10 CV. The NS1 and NS1-ZF5.3 proteins were not stable for overnight dialysis and were therefore either immediately labeled with rhodamine (see Rhodamine Labeling of NS1 Proteins below) or buffer-exchanged into Storage Buffer 3 using a PD-10 desalting column. Protein concentration was determined using absorbance at 280 nm as described above. LC/MS was used to confirm the identity of all proteins, which were then stored at -80°C until use.

**Sortase-mediated rhodamine labeling of DHFR proteins**

$DHFR^{Rho}$ and ZF5.3-$DHFR^{Rho}$ were generated by reaction with a $GGGK^{Rho}$ peptide and either the StrepTagII-SrtA7m-His6 enzyme (for $DHFR^{Rho}$) or the His6-SUMO-SrtA7m enzyme with the His6 tag cleaved (for ZF5.3-$DHFR^{Rho}$). The different enzymes were chosen for their ability to be separated from the reaction mixture, as described below. The $GGGK^{Rho}$ peptide was synthesized as previously described (2). For labeling, DHFR and ZF5.3-DHFR were diluted to 35 µM in a 1.5 mL solution containing 75 µM of the appropriate SrtA7m enzyme and 200 µM $GGGK^{Rho}$ in 20 mM Tris, 150 mM KCl, 1 mM TCEP pH 7.5. The reaction was incubated with gentle rotation for 4 h at 4°C.

After 4 h, $DHFR^{Rho}$ was separated from the reaction mixture by incubating with 1 mL TALON® resin for 1 h at 4°C and applying the solution to a gravity column. The gravity column was washed with 10 mL of Wash Buffer 2. Because the His6 tag is C-terminal to the LPETGG motif on DHFR, successful reaction with sortase should result in loss of the His6 tag and therefore $DHFR^{Rho}$ is collected in the column washes and flowthrough. Fractions containing fluorescent DHFR were analyzed by SDS-PAGE using a fluorescence imager, pooled, and desalted into Storage Buffer 1 using a disposable PD-10 desalting column (Cytiva) to remove excess $GGGK^{Rho}$. The final rhodamine-labeled protein was concentrated using a 10 kDA MWCO Amicon spin concentrator.

To separate ZF5.3-$DHFR^{Rho}$ from the other reaction components, the reaction mixture was incubated with 1 mL TALON® resin for 1 h at 4°C and applied to a gravity column. In this case, although successful reaction with sortase will still result in loss of the His6 tag, ZF5.3-DHFR binds the TALON® resin due to nonspecific interactions between ZF5.3 and the resin. The resin was washed with 10 mL Wash Buffer 2 to remove excess $GGGK^{Rho}$ and SrtA7m. ZF5.3-$DHFR^{Rho}$ was eluted with 15 mL Elution Buffer 1 and dialyzed overnight at 4°C into Storage Buffer 1 + 100 µM $ZnCl_2$. After dialysis, the protein was concentrated using a 10 kDA MWCO Amicon spin concentrator.

For $DHFR^{Rho}$ and ZF5.3-$DHFR^{Rho}$, the identities of the purified proteins were confirmed using LC/MS. Protein concentrations were calculated using the Pierce™ 660 nm Protein Assay. The concentrations of rhodamine-labeled proteins were determined by calculating the rhodamine concentration in each sample using the Beer-Lambert law $A = \varepsilon bC$ where A = the absorbance at 570 nm on a NanoDrop spectrophotometer, $\varepsilon$ = the extinction coefficient of lissamine rhodamine B in water (112000 $M^{-1}$ $cm^{-1}$), b = the path length in cm, and C = the concentration in M. The labeling efficiency was calculated by dividing the rhodamine concentration by the concentration of total protein. All proteins were stored at -80°C until use.

**Rhodamine labeling of SNAP-tag proteins**

All SNAP proteins were tagged with rhodamine using a self-labeling reaction between the SNAP-tag variant and a rhodamine dye functionalized with a benzylguanine moiety (BG-Rho). BG-rho was synthesized as previously described (1). BG-amine (3 mg, 11 µmol) was dissolved in 500 µL anhydrous DMSO and added to 3 equivalents of lissamine rhodamine B sulfonyl chloride (33 µmol) and 6 equivalents of N,N-Diisopropylethylamine (66 µmol). The reaction was stirred under nitrogen at room temperature overnight. The next day, BG-Rho was purified with a reverse-phase preparative HPLC (Waters Prep 150 System) using a C18 column (CSH C18 19 x 150 mm OBD

Column 5 μm), and fractions with the desired product were identified using LC/MS. These fractions were pooled, lyophilized, and resuspended in DMSO for labeling experiments. The concentration of BG-rho was determined using absorbance at 570 nm as described above.

$SNAP^{Rho}$, $AGT54^{Rho}$, and GE-$AGT^{Rho}$ were generated by incubating a 40 μM solution of each respective protein with 80 μM BG-Rho in Labeling Reaction Buffer (20 mM Tris, 150 mM NaCl, 10% glycerol pH 7.5) overnight with rotation at 4°C. ZF5.3-$SNAP^{Rho}$, ZF5.3-$AGT54^{Rho}$, and ZF5.3-GE-$AGT^{Rho}$ were generated by incubating a 20 μM solution of each respective protein with 40 μM BG-Rho in Reaction Buffer for 24 h at 4°C with rotation. Reaction progress in all cases was monitored by LC/MS until completion. Excess BG-Rho was removed from the reactions using a PD-10 desalting column and the rhodamine-labeled proteins were concentrated using a 65% w/v ammonium sulfate precipitation as described above. The $SNAP^{Rho}$, $AGT54^{Rho}$, and GE-$AGT^{Rho}$ protein pellets were resuspended in Storage Buffer 2, while the ZF5.3-$SNAP^{Rho}$, ZF5.3-$AGT54^{Rho}$, and ZF5.3-GE-$AGT^{Rho}$ pellets were resuspended in Storage Buffer 2 + 100 μM $ZnCl_2$. The rhodamine-labeled protein concentrations were determined using absorbance at 570 nm as described above, with the extinction coefficient (ε) for lissamine rhodamine B measured in salty storage buffer (64417 $M^{-1}$ $cm^{-1}$). All proteins were stored at -80°C until use.

**<u>Rhodamine labeling of NS1 proteins</u>**

Fractions containing pure NS1 or NS1-ZF5.3 from the Strep affinity purification were pooled and immediately incubated with 5 equiv of lissamine rhodamine B-maleimide (Tenova) in the buffer with which they were eluted off the column. The solution was incubated with rotation at RT for 1 h. To remove excess dye, the protein was desalted using a PD-10 column and eluted into Storage Buffer 3. The rhodamine-labeled protein concentrations were determined by absorbance at 570 nm. Concentration of total protein was determined using absorbance at 280 nm. Labeling efficiencies were determined by dividing the concentration of rhodamine-labeled protein by the concentration of total protein and ranged between 22-53%. LC/MS was used to confirm the identity of all proteins, which were then stored at -80°C until use.

**<u>Circular dichroism</u>**

Circular dichroism (CD) measurements were performed and analyzed using existing protocols (8). All spectra were recorded using either an AVIV Biomedical, Inc. (Lakewood, NJ) Circular Dichroism Spectrometer Model 410 or a JASCO J-1500 Circular Dichroism Spectropolarimeter. In all cases, spectra were recorded using a 0.1 cm pathlength quartz cuvette (Starna). Wavelength-dependent CD spectra were collected between 300 and 200 nm in 1 nm intervals with an averaging time of 5 sec and normalized to a baseline buffer-only spectrum. Temperature-dependent CD spectra were collected between 5 and 90°C measuring every 2° with an equilibration time of 120 s at each temperature step. The wavelength used for the temperature melt for each protein was chosen based on the wavelength spectra and set to 210 nm for DHFR proteins, 222 nm for SNAP-tag proteins, and 218 nm for NS1 proteins. For DHFR and ZF5.3-DHFR, all spectra were collected at 20 μM protein concentration in 25 mM Tris, 150 mM KCl, 1 mM TCEP pH 7.5 (+25 μM $ZnCl_2$ for ZF5.3-DHFR only). For measurements with methotrexate (MTX), DHFR or ZF5.3-DHFR was pre-incubated with one equiv of MTX for 30 min prior to measurements. For $SNAP^{Rho}$, $AGT54^{Rho}$, GE-$AGT^{Rho}$, ZF5.3-$SNAP^{Rho}$, ZF5.3-$AGT54^{Rho}$, and ZF5.3-GE-$AGT^{Rho}$, all spectra were collected at 12 - 20 μM protein concentration in 25 mM Tris,

150 mM NaCl, 1 mM TCEP pH 7.5 (+25 µM $ZnCl_2$ for ZF5.3 variants). For NS1 and NS1-ZF5.3, the spectra were collected at 18 µM in a buffer composed of 20 mM Tris, 150 mM KCl, 0.5 mM TCEP pH 7.5. Raw ellipticity values were converted to mean residue ellipticity using eq (1):

$$\theta = \frac{m^\circ \bullet MRW}{l \cdot C} \qquad (1)$$

where θ = mean residue ellipticity in deg • $cm^2$ • $dmol^{-1}$, m° = raw ellipticity in millidegrees, MRW = mean residue weight (calculated as the molecular weight divided by the number of backbone amide bonds), $l$ = path length of the cuvette in millimeters, and C = concentration of the protein in mg $mL^{-1}$. For temperature melts, normalized intensity was calculated by normalizing all ellipticity values to the value at 5°C.

**DHFR activity assay:**

The catalytic activities of purified DHFR and ZF5.3-DHFR were assessed using a commercially available Dihydrofolate Reductase Assay kit (Sigma). The assay monitors the NADPH-dependent reduction of dihydrofolate (DHFA) to tetrahydrofolate (THFA) by measuring the loss in absorbance at 340 nm over time using a UV/Vis spectrometer (Cary 60 UV-Vis, Agilent Technologies). Purified DHFR or ZF5.3-DHFR was added to a final concentration of 50 nM in a 1 mL solution of 1x activity assay buffer provided in the kit. The solution was transferred to a 1 cm quartz cuvette, followed by addition of 6.6 µL of 10 mM NADPH. The cuvette was inverted to mix the reaction components. In quick succession, 5 µL of 10 mM DHFA was added, the mixture was inverted again, and the reaction progress was monitored by absorbance at 340 nm. Absorbance readings were taken every 15 s for 150 s. For measurements in the presence of MTX, MTX was added to a final concentration of 100 nM and incubated with the protein for 5 min before addition of NADPH and DHFA.

The specific activity (in units per mg) of each enzyme was calculated using eq (2):

$$Units/mg = \frac{(\Delta OD/min(sample) - \Delta OD/min(blank)) \cdot d}{12.3 \cdot V \cdot mg/mL} \qquad (2)$$

where ΔOD/min(sample) = the slope of the 340 nm absorbance curve for the sample, ΔOD/min(blank) = the slope of the 340 nm absorbance curve for a buffer blank, d = dilution factor of the enzyme sample, 12.3 = the extinction coefficient ($mM^{-1}$ $cm^{-1}$) for the DHFR reaction at 340 nm, V = the volume of enzyme used in the reaction, and mg/mL = concentration of the DHFR protein in mg $mL^{-1}$.

**Cell culture**

Saos-2 cell stocks were purchased from the UC Berkeley Cell Culture Facility. Saos-2 cells were cultured in McCoy's 5A medium with phenol red containing 15% fetal bovine serum (FBS), penicillin and streptomycin (P/S, 100 units/mL and 100 µg/mL, respectively), 1 mM sodium pyruvate, and 2 mM GlutaMax.

#### Cellular workup for delivery experiments

All fluorescence-based intracellular delivery experiments (confocal microscopy, flow cytometry, and fluorescence correlation spectroscopy) followed previously published protocols (9). The day prior to experiments, $0.2 \cdot 10^6$ Saos-2 cells were plated in a 6-well dish in McCoy's 5A medium +15% FBS, without phenol red or P/S. The following day, cells were incubated with a 0.5 mL (SNAP proteins) or 1 mL (DHFR proteins and NS1 proteins) solution of rhodamine-labeled protein diluted to the appropriate concentration (0.1 - 2 µM) in clear McCoy's 5A medium without FBS or P/S. During the incubation, a fibronectin solution diluted into DPBS (0.1 mg/mL) was added to the appropriate number of wells of an 8-well microscopy dish. In a separate well of the same dish, 200 µL of a 100 nM solution of AlexaFluor 594 hydrazide dye in water was added for FCS analysis to calculate the microscope focal volume. The dish was incubated at 37°C, 5% $CO_2$ for the duration of the cell incubation and work-up. During the last 5 min of the cell incubation (0.5 - 2 h total length), Hoechst 33342 nuclear dye was added to each well of the 6-well dish at a final concentration of 300 nM. After the incubation was over, the protein-containing solution was removed and the cells were washed three times with 2 mL of DPBS per wash. To remove exogenously bound protein, cells were trypsinized with 500 µL trypsin (no phenol red). The trypsin reaction was quenched with 1 mL clear McCoy's 5A medium + 15% FBS and each well was thoroughly washed with another 1 mL of the same media. The cells were pelleted in a 15 mL Falcon tube and washed one time with 1 mL of clear DMEM + Hepes. During this centrifugation step, the microscopy dish was removed from the incubator and fibronectin-containing wells were washed three times with 200 uL DPBS per wash. The DPBS was left in each well until cells were ready to add. The pelleted cells were then resuspended in 600 µL DMEM + Hepes. Of this, 300 µL was added to each fibronectin-coated well of the microscopy dish for confocal microscopy and FCS analysis, and transferred to 37°C, 5% $CO_2$ to allow the cells to adhere flatly. The remaining 300 µL of cells were pelleted, resuspended in 200 µL DPBS, and transferred to a 1.5-mL Eppendorf tube for flow cytometry.

#### Flow Cytometry

For each sample processed as described above, 10,000 cells were analyzed using flow cytometry. Forward scatter and side scatter gating parameters were established to select only whole cells using the same settings as reported previously (2, 3). For all experiments, >75% of cells were included in the gated population. The fluorescence intensities for Hoechst 33342 (excited using the violet laser at 405 nm, detected emission of 440 ± 50 nm) and lissamine rhodamine B (excited using the yellow laser at 561 nm, detected emission of 586 ± 16 nm) channels were monitored using an Attune NxT Focusing Flow Cytometer. The median fluorescence intensity (MFI) for the lissamine rhodamine B channel was reported for the gated population of cells, corrected for the rhodamine labeling efficiency of each protein when appropriate. Each flow cytometry experiment was performed with a minimum of two biological replicates (**Table S3**).

#### Confocal microscopy and fluorescence correlation spectroscopy (FCS)

The procedures used for confocal microscopy and FCS have been described previously (3, 9). Experiments were performed with a STELLARIS 8 microscope (Leica Microsystems) with a Leica DMi8 CS scanhead, a HC Plan-Apo 100/1.4NA STED white oil immersion objective (used for

STED imaging), a HC Plan-Apo 63x/1.4NA water immersion objective (used for FCS measurements), a pulsed white-light laser (440 nm- 790 nm; 440 nm: > 1.1 mW; 488 nm: > 1.6 mW; 560 nm: > 2.0 mW; 630 nm: > 2.6 mW; 790 nm: > 3.5 mW, 78 MHz), and a pulsed 775nm STED laser. All confocal imaging was performed using HyD S or HyD X detectors in analog mode, while FCS measurements were carried out using only a Hybrid HyD X detector. All microscopy experiments were performed at 37°C (monitored using Oko-Touch) and 5% $CO_2$ in a blacked out cage enclosure from Okolab. Before the start of each experiment, the correction collar of the objective was adjusted by maximizing the counts per molecule for the AlexaFluor 594 hydrazide dye standard; minor fluctuations in the correction collar are expected based on the variable thickness of the glass-bottom microscopy dishes (LabTek™). Lissamine rhodamine B was excited at 561 nm with a fluorescence filter of 570 - 660 nm, and the pinhole S5 of the laser was set to 1 AU. Before beginning *in cellulo* experiments, ten five-second autocorrelation traces were obtained using the well containing 100 nM AlexaFluor 594 hydrazide dye standard to calculate the focal volume of the microscope; see below. For *in cellulo* data procurement, a confocal microscopy image of the cells was used to position the crosshairs of the microscope laser in the cytosol of cells within the frame. Areas with punctate fluorescence indicative of endosomally entrapped protein were avoided. All FCS measurements consisted of ten five-second traces. A minimum of 35 cells per condition were measured for each biological replicate, and a minimum of two biological replicates were collected for each condition. Expected diffusion times ($\tau_{\text{diff}}$) for rhodamine-labeled DHFR, SNAP, and NS1 proteins were obtained by measuring *in vitro* autocorrelation traces for 50-200 nM solutions of each protein in DMEM media (25 mM Hepes, no phenol red) at 37°C.

**<u>Analysis of FCS data</u>**

Autocorrelation traces obtained from FCS measurements were analyzed using a custom MATLAB script (2, 3, 9). To extract quantitative information from *in cellulo* data, the effective confocal volume of the microscope must be known. This value was determined using eqs 3-5 by and the *in vitro* autocorrelation traces for the AlexaFluor 594 hydrazide standard measured at the start of each experiment, which has a known diffusion coefficient in water (10). These traces were fitted to a 3D diffusion equation (eq 3):

$$G(\tau) = \frac{1}{N} \cdot \frac{1}{(1+\frac{\tau}{\tau diff})\sqrt{1+(s^2\frac{\tau}{\tau diff})}} \qquad (3)$$

where N = the average number of molecules detected in the focal volume ($V_{\text{eff}}$), $\tau_{\text{diff}}$ = the average diffusion time that a molecule requires to cross $V_{\text{eff}}$, and s = the structure factor (the ratio of the radial to axial dimensions of the focal volume). The structure factor was measured to be 0.17 using the autocorrelation function of AlexaFluor 594 in water at 25°C and fixed for all subsequent analysis. $V_{\text{eff}}$ can be extracted from these data by inserting the $\tau_{\text{diff}}$ value derived from eq (3) to calculate $\omega_1$ in eq (4):

$$\omega_1 = \sqrt{4 \cdot D \cdot \tau_{diff}} \qquad (4)$$

where $\omega_1$ = the lateral extension of the confocal volume and D = the known diffusion coefficient of AlexaFluor 594 in water at 37°C ($5.20 \times 10^{-6}$ $cm^2\,s^{-1}$). $V_{eff}$ can then be directly calculated from eq (5):

$$V_{eff} = \pi^{\frac{3}{2}} \cdot \omega_1 \cdot \frac{1}{s} \qquad (5)$$

The average $V_{eff}$ for all experiments ranged from 0.25-0.4 fL.

Autocorrelation traces derived from *in cellulo* measurements were fitted using a 3D anomalous diffusion equation (eq 6):

$$G(\tau) = \frac{1}{N} \cdot \frac{1}{(1+\frac{\tau}{\tau diff})^{\alpha} \sqrt{1+s^2(\frac{\tau}{\tau diff})}^{\alpha}} + G(\infty) \qquad (6)$$

where N = the average number of molecules in the focal volume, $\tau_{diff}$ = the average diffusion time that a molecule requires to cross $V_{eff}$, α = the anomalous diffusion coefficient, and s = the structure factor (0.17). Parameters resulting from the fitting (including diffusion time, anomalous diffusion coefficient, and counts per molecule) were filtered as reported previously (11), and only fitted curves that passed this filtering and displayed Chi-squared values under 40 (for DHFR proteins) or 100 (for SNAP and NS1 proteins) were selected for further analysis. Expected $\tau_{diff}$ values for *in cellulo* samples were predicted from autocorrelation traces for *in vitro* protein samples in DMEM that were fitted using eq (3). Only curves that displayed *in cellulo* $\tau_{diff}$ values within 2-15-fold of the *in vitro* value were selected. Shorter diffusion times can indicate degradation, while longer diffusion times may reflect aggregated material with abnormal diffusion dynamics. The concentration (C) of protein in the cytosol was then calculated using the value of N derived above in eq (7):

$$C = \frac{N}{N_A \cdot V_{eff}} \qquad (7)$$

where $N_A$ = Avogadro's number ($6.023 \times 10^{23}$ $mol^{-1}$). When relevant, the concentration values were adjusted for fluorophore labeling efficiency. At least 15 concentration values from curves that pass all filters were used for each FCS condition (**Table S3**).

#### Cytosolic fractionation

The day before the experiment, $5.0 \times 10^6$ Saos-2 cells were plated in a 100 mm dish in McCoy's 5A medium + 15% FBS (no phenol red, no P/S). The following morning, the cells were washed three times with DPBS and the media was replaced with 10 mL of McCoy's 5A media (no FBS, no phenol red, no P/S) containing a 1 µM solution of DHFR or ZF5.3-DHFR. The cells were incubated at 37°C, 5% $CO_2$ for 1 hour. After incubation, the cells were washed three times with DPBS and lifted from the dish with trypsin. The trypsin reaction was quenched with 15 mL of McCoy's 5A media + 15% FBS (no phenol red, no P/S) and the cells were transferred to a 50 mL conical tube. The dish was washed with an additional 5 mL of the same media. The cells were pelleted by centrifugation at 200 g for 3 min and washed by resuspension with 3 mL of DPBS. After centrifugation, the cells were washed by resuspending in 1 mL of precooled buffered isotonic sucrose (290 mM sucrose, 10 mM imidazole pH 7.0, 1 mM DTT, and 1 cOmplete protease inhibitor tablet) and pelleted again. The cells were then resuspended in 100 µL of the isotonic sucrose buffer and transferred to 0.5 mL tubes with 1.4 mm ceramic beads (Omni International). A Bead Ruptor was used to lyse the cells for 10 sec at speed 1. Homogenized cells were then transferred to polycarbonate ultracentrifuge tubes and balanced within 1 mg. The cytosolic fraction was separated from the membrane fraction by centrifugation at 350 kg for 1 h at 4°C (Beckman Coulter, TLA-100 rotor). The supernatant (cytosol) was immediately removed and transferred to a 1.5 mL Eppendorf tube. The pellet (containing endosomes) was resuspended in 60 µL of 8 M urea and boiled at 95°C for 5 min to fully dissolve the pellet. Samples for Western blot analysis were prepared by mixing 20 µL of cytosolic or endosomal membrane fractions with 5 µL of 5x SDS gel loading dye and boiling at 95°C for 5 min. Western blot analysis was performed using primary antibodies against His-tag (CST #2365), tubulin (CST #2125), and Lamp1 (CST #9091) and a secondary HRP-linked anti-Rabbit IgG antibody (CST #7074).

#### siRNA knockdowns

Knockdowns for VPS39 and TGF-BRAP1 were evaluated using the siRNAs listed in **Table S1** and were performed as described previously (10). Four days before delivery, 0.085 × $10^6$ Saos-2 cells were plated in a 6-well dish in McCoy's 5A media +15% FBS (no phenol red, no P/S). The next day, cells were transfected with the siRNA of interest at a final concentration of 100 nM using Lipofectamine RNAi/MAX transfection agent in McCoy's 5A media + 15% FBS (no phenol red, no P/S) and incubated for 4 h at 37°C, 5% $CO_2$. Cells were then washed three times with DPBS and incubated for 72 h at 37°C, 5% $CO_2$ in McCoy's 5A media + 15% FBS (2 mL/well, no phenol red, no P/S) before delivery experiments.

#### RT-qPCR analysis of siRNA knockdowns

RT-qPCR was used to confirm siRNA-mediated knockdown of the desired genes after 72 h (10). cDNA extracted from Saos-2 cells was amplified with gene-specific primers (150 nM, **Table S1**) using SsoFast EvaGreen Supermix (Bio-Rad) on a Bio-Rad CFX96 real-time PCR detection system. Three biological replicates were performed for each knockdown, and each sample was run in triplicate. Each RT-qPCR analysis included the gene that was knocked down, GAPDH as a reference gene, and negative controls lacking either reverse transcriptase or template cDNA.

#### Analysis of ZF5.3-DHFR$^{Rho}$ localization in Lamp1+ vesicles using confocal microscopy and STED

Two days prior to experiments, 0.05 × $10^6$ Saos-2 cells were plated in a 4-well glass-bottom µslide microscopy dish (Ibidi) in McCoy's 5A media + 15% FBS (no phenol red, no P/S). The next day, cells were transfected with a plasmid encoding Lamp1-HaloTag (Addgene) using FuGENE HD transfection reagent (Promega) according to the manufacturer's protocol. After 6 h, the cells were washed 2 times with DPBS and incubated overnight in McCoy's 5A media + 15% FBS (no phenol red, no P/S) at 37°C, 5% $CO_2$. On the day of the experiment, cells were incubated with a chloroalkane-modified silicon rhodamine dye (SiR-CA) (12) for 1 h at a final concentration of 2 µM in OptiMEM (700 µL final volume). After this incubation, the cells were washed twice with DPBS and the media was replaced with McCoy's 5A media (no FBS, no phenol red, no P/S) containing 0.5 µM ZF5.3-DHFR$^{Rho}$ for 1 hr. The cells were then washed three times with DPBS, trypsinized to remove exogenous protein, and replated in a fibronectin-coated 4-well glass-bottom µslide microscopy dish in DMEM (+Hepes). The sample was then imaged on a Leica Stellaris 8 STED-capable confocal microscope with a HC PL APO CS2 100x/1.4 NA Oil Leica STED-White objective. Beam path settings: ZF5.3DHFR$^{Rho}$ (Excited at 573 nm with 0.5% laser power, HyD X detector window: 579 nm - 636 nm), SiR-CA (excited at 652 nm with 20% laser power, HyD X detector window: 662 nm - 750 nm); depletion at 775 nm at 30% laser power. Detectors were in photon counting mode and were set up to acquire signal simultaneously. Pinhole was set to 1 AU at 580 nm emission. Scanhead settings: Format: 1024 x 1024; pixel size: 28 nm pixel size; pixel dwell time: 3.8375 **µ**s. TauSTED parameters were adjusted in post-processing. Images were processed using FIJI software package. Images were cropped and a smoothening filter was applied before plotting line profiles.

#### Statistics

All statistical tests were performed using GraphPad Prism 9 software and are detailed in the appropriate figure legends. Flow cytometry and FCS data were analyzed using Brown-Forsythe and Welch ANOVA statistical tests followed by unpaired t-tests with Welch's correction.

#### Data Availability

The data in this study are available from the corresponding author upon request. The MATLAB® script for FCS analysis is available from GitHub (https://github.com/schepartzlab/FCS).

#### Supplementary Figures

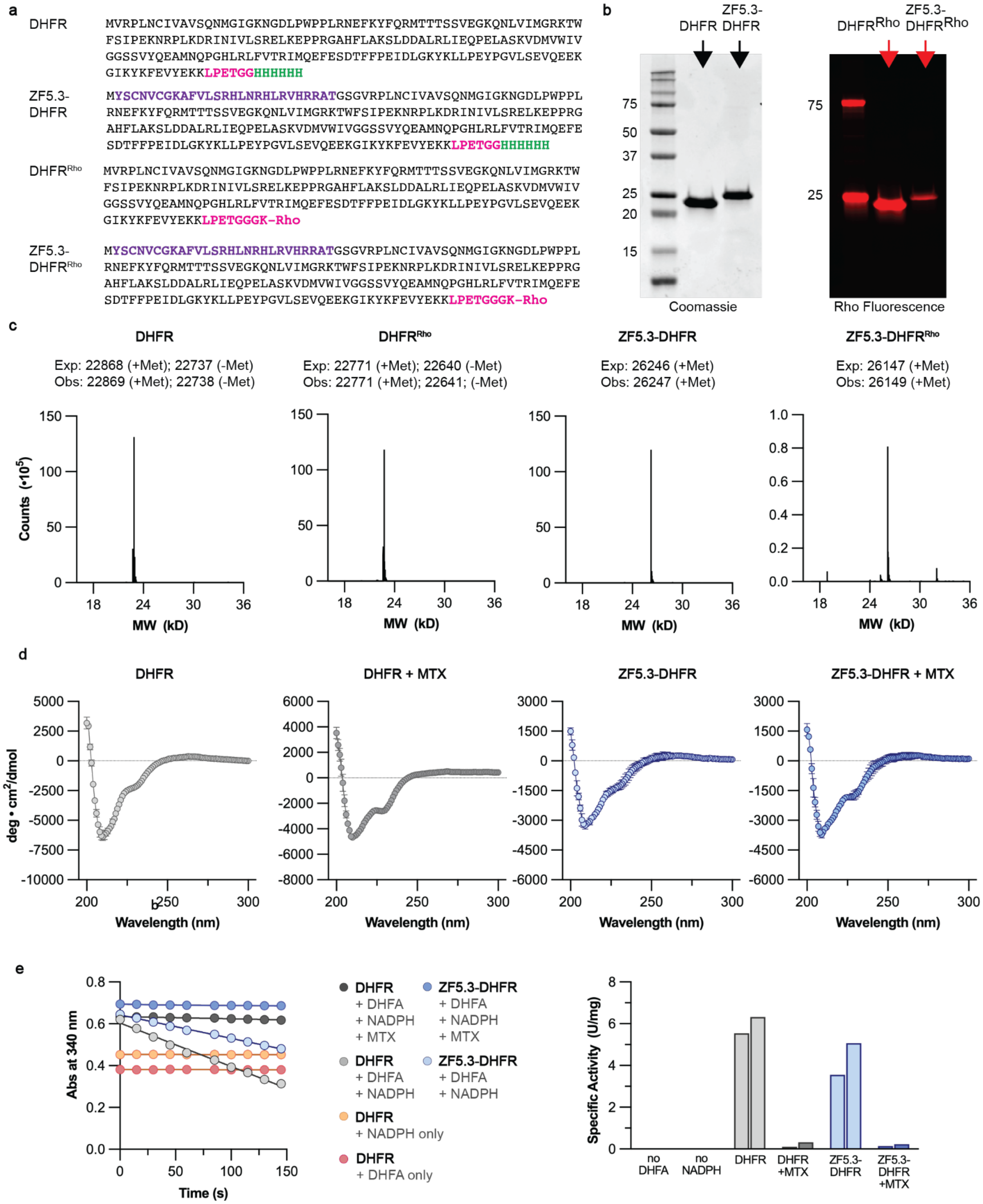


**Fig. S1: Purification and characterization of DHFR, ZF5.3-DHFR, and rhodamine-tagged variants. a,** DHFR proteins used in this study comprised the complete sequence of murine DHFR with or without a ZF5.3 module (purple, 27 aa) appended to the N-terminus. All proteins also contained a His6 tag for affinity purification (green) and a C-terminal LPETGG sortase recognition motif (magenta) to enable subsequent labeling with rhodamine, as described previously (1–3) and in Methods. **b,** SDS-PAGE analysis of purified DHFR, ZF5.3-DHFR, $DHFR^{Rho}$ and ZF5.3-$DHFR^{Rho}$, with proteins visualized either using Coomassie stain or fluorescence imaging. Labeling efficiency ranged between 9 and 88%. **c,** LC/MS analysis of all DHFR proteins confirms the correct identity of each. For DHFR and $DHFR^{Rho}$, prominent peaks were observed for the protein +/- the N-terminal methionine residue. **d,** Wavelength-dependent circular dichroism (CD) analysis of DHFR and ZF5.3-DHFR (20 μM) at room temperature in a buffer composed of 25 mM Tris, 150 mM KCl, pH 7.2 and in the presence and absence of 1 equivalent methotrexate (MTX). Spectra were collected between 200 and 300 nm at 1 nm intervals with an averaging time of 5 seconds. Data shown are from at least two biological replicates and are represented as mean ± SEM. **e,** All purified DHFR proteins retain catalytic activity. DHFR catalyzes the reduction of dihydrofolate (DHFA) to tetrahydrofolate (THFA) using NADPH as an electron donor. Plot on left illustrates the time-dependent loss in NADPH absorbance at 340 nm of solutions containing the indicated components. Conversion of DHFA to THFA occurs only when all reaction components are present; activity is abolished when 1) MTX is present (dark gray and dark blue), 2) DHFA is missing (orange), or 3) NADPH is missing (pink). Absorbance plot on left is representative of two activity assays performed on separate batches of protein. Plot on right shows the specific activity calculated in units per mg of protein for the same six conditions. Each bar represents activity from a separate batch of purified protein. One unit will convert 1.0 μmol of DHFA to THFA in 1 minute at pH 7.5 at room temperature.

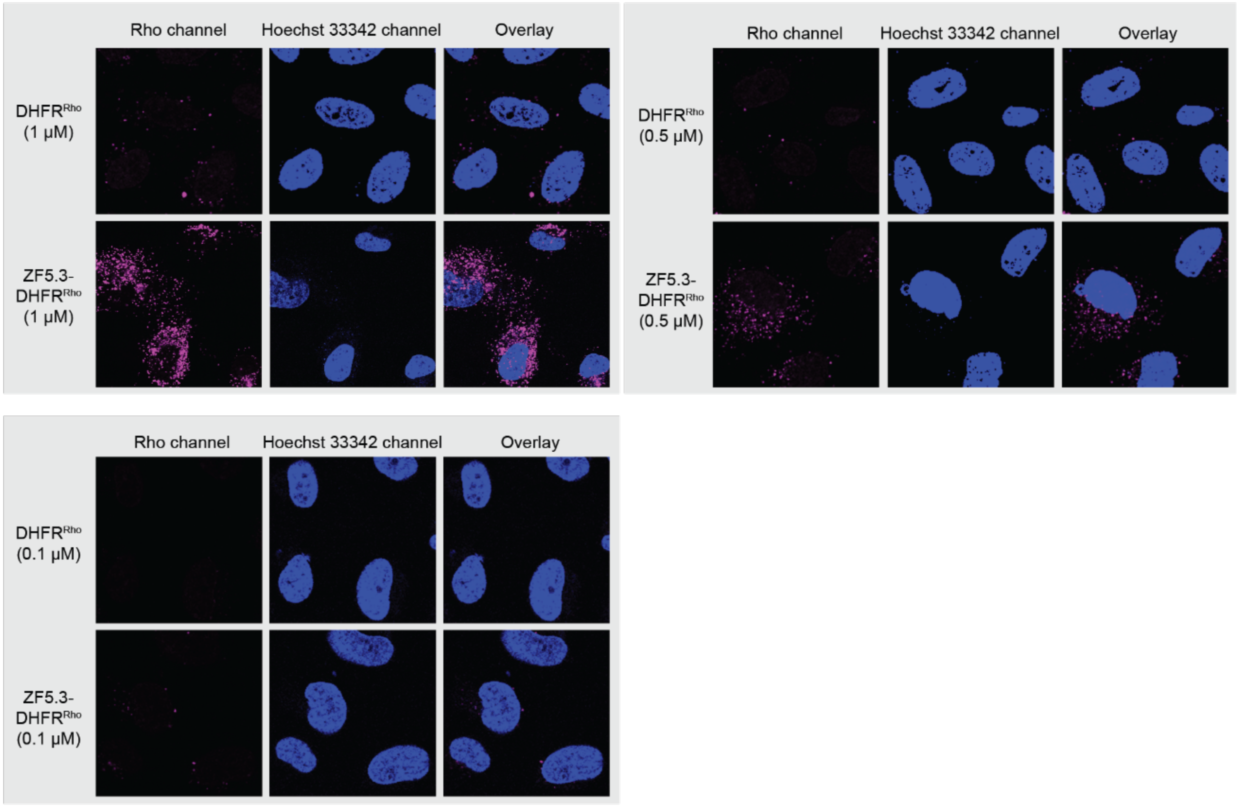


**Fig. S2: 2D confocal microscopy images depicting total intracellular fluorescence of Saos-2 cells treated with DHFR$^{Rho}$ and ZF5.3-DHFR$^{Rho}$.** Saos-2 cells were incubated with the given concentration of DHFR$^{Rho}$ or ZF5.3-DHFR$^{Rho}$ for 1 h, followed by three DPBS washes and trypsinization to remove exogenous protein. Cells were then replated in a fibronectin-coated 8-well microscopy dish and visualized using confocal microscopy. Nuclear fluorescence was detected by adding 300 nM Hoechst 33342 for the final 5 minutes of the protein incubation period. The results shown are representative of at least two biological replicates.

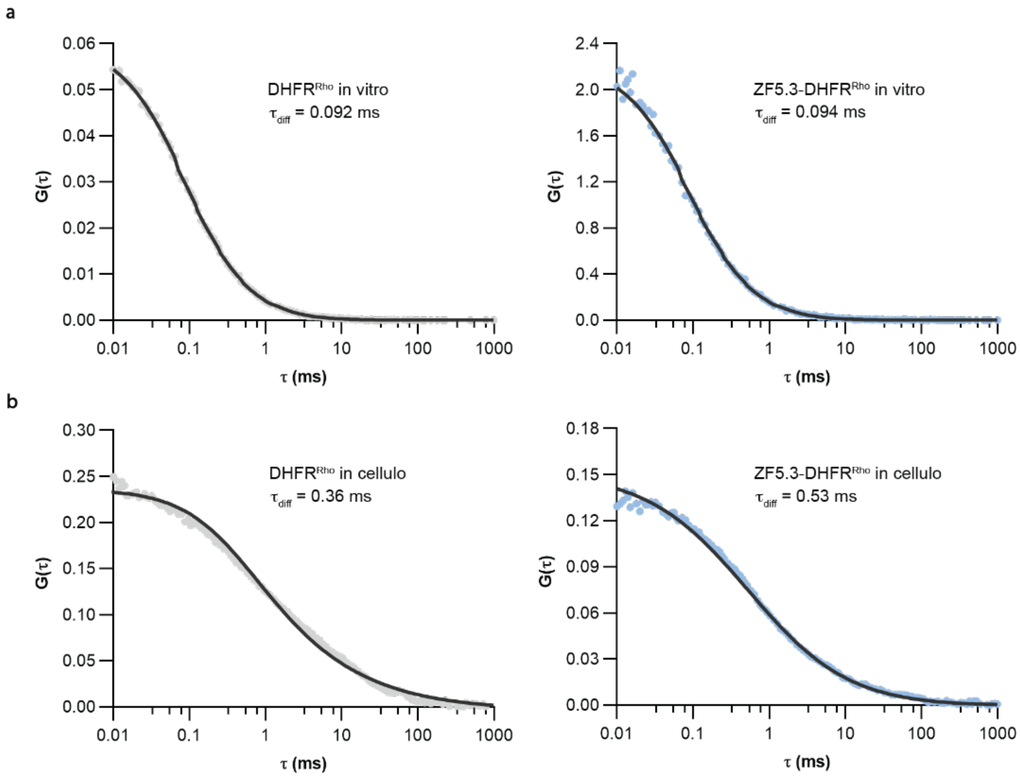


**Fig. S3. Autocorrelation traces generated during FCS analysis of DHFR$^{Rho}$ and ZF5.3-DHFR$^{Rho}$ *in vitro* and in Saos-2 cells. a,** Autocorrelation traces generated for *in vitro* samples of 100-500 nM DHFR$^{Rho}$ and ZF5.3-DHFR$^{Rho}$ in DMEM cell media are shown. All traces were fitted to a 3D diffusion model (solid dark curve) to obtain *in vitro* diffusion times ($t_{diff}$) as described in Methods. **b,** Representative autocorrelation traces of Saos-2 cells treated with 500 nM DHFR$^{Rho}$ and ZF5.3-DHFR$^{Rho}$ for 1 h. Traces were fitted using a 3D anomalous diffusion model as previously described[3] to obtain *in cellulo* diffusion times. *In cellulo* diffusion times are longer than those *in vitro* due to increased viscosity and impaired diffusion in the cytosol.

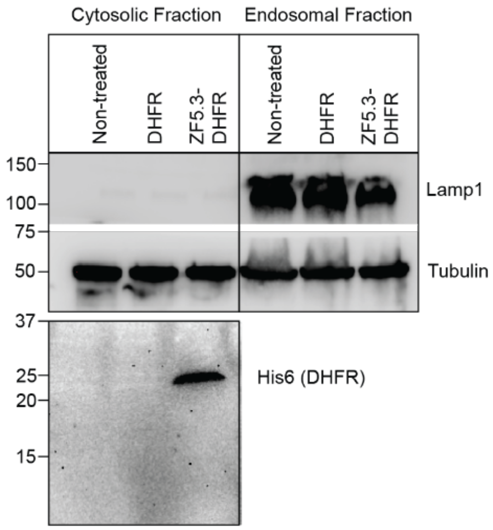


**Fig. S4: Cytosolic fractionation of cells treated with DHFR and ZF5.3-DHFR.** Western blot analysis was performed for cytosolic or endosomal fractions isolated from Saos-2 cells treated with clear McCoy's media alone (non-treated) or 1 μM DHFR or ZF5.3-DHFR for 1 hour. After treatment, cells were lysed and the cytosol was isolated by ultracentrifugation at 350 kg for 1 h; see Methods for detailed cytosolic fractionation procedure. Western blot was performed with antibodies against endocytic (Lamp1) and cytosolic (tubulin) markers for both the endosomal and cytosolic fractions. Gel results are representative of two biological replicates.

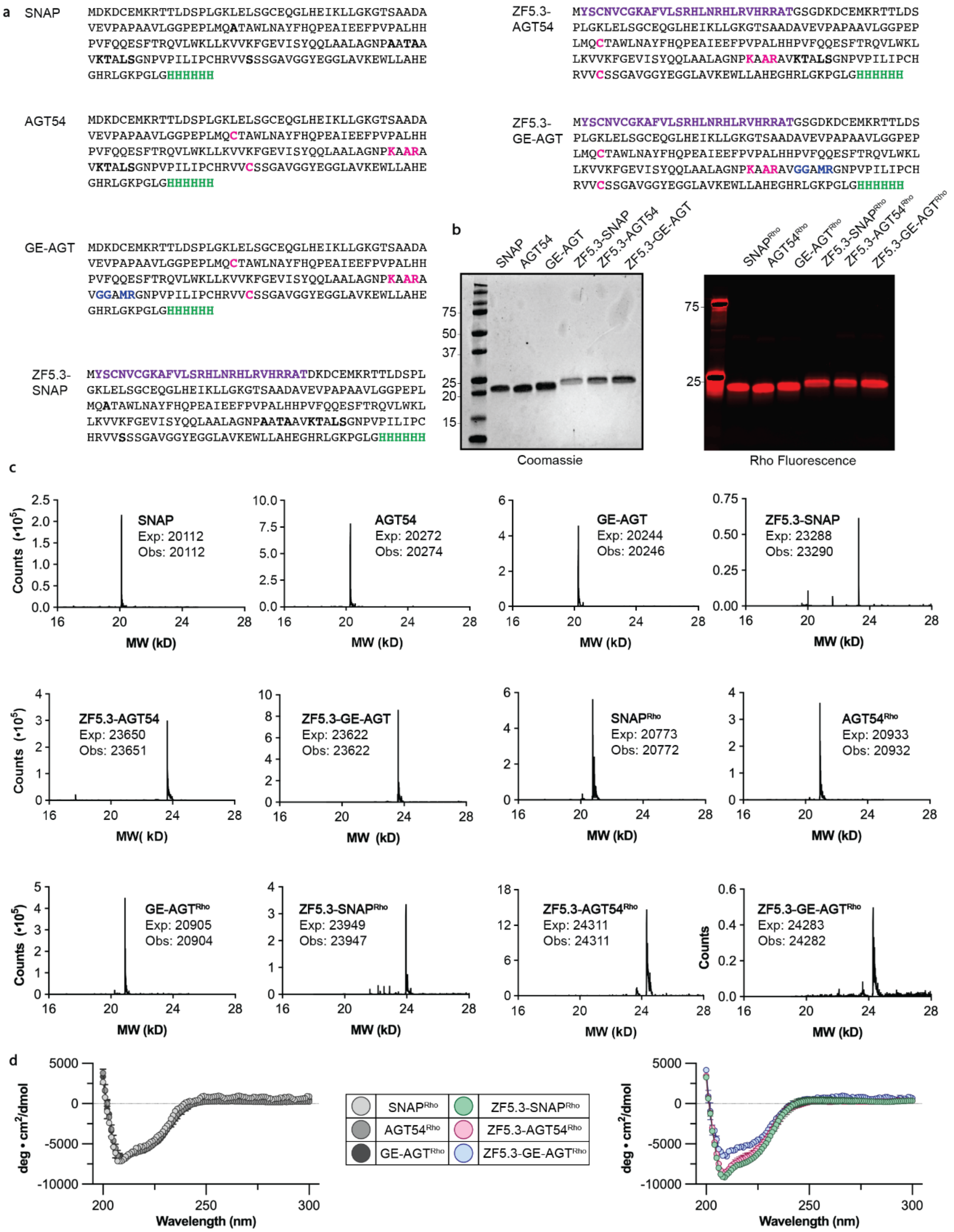


**Fig. S5: Purification and characterization of SNAP variants via SDS-PAGE, LC/MS, and circular dichroism. a,** The SNAP sequences chosen for this study were adapted from the literature (4) and modified with an N-terminal ZF5.3 moiety (purple) and a C-terminal His6 tag for affinity purification (green). Mutations introduced to AGT54 and GE-AGT to decrease thermostability are shown in magenta; mutations introduced only to GE-AGT are shown in blue. All proteins were reacted with a lissamine rhodamine B dye derivatized with benzylguanine to enable fluorescent labeling, as previously described (1). **b**, SDS-PAGE analysis of all purified variants with or without an attached rhodamine dye were visualized using coomassie stain (for unlabeled proteins) or fluorescence imaging (for rhodamine-modified proteins). **c**, LC/MS analysis confirmed the identity of all unlabeled and rhodamine-labeled proteins. Additional peaks for some samples with successive +98 Da molecular weights are observed due to sulfate adducts formed during ammonium sulfate precipitation. **d,** Wavelength- dependent CD analysis of rho-labeled SNAP and ZF5.3-SNAP variants (20 μM) at room temperature in a buffer composed of 20 mM Tris, 150 mM NaCl pH 7.5. Measurements were recorded between 200 and 300 nm in 1 nm intervals with an averaging time of 5 seconds. Data shown are from at least two biological replicates and are represented as mean ± SEM.

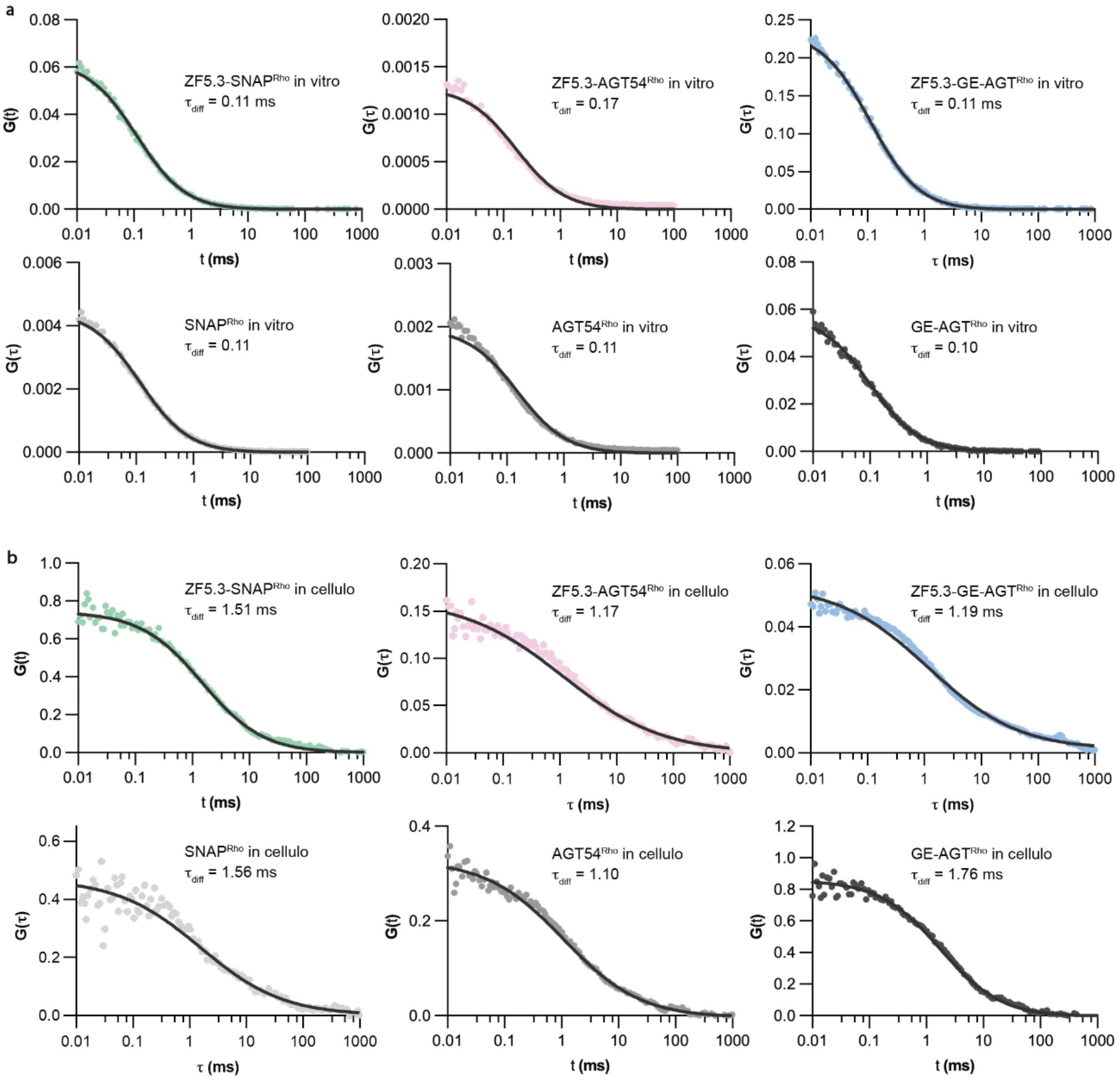


**Fig. S6: *In vitro* and *in cellulo* autocorrelation traces for all SNAP$^{\text{Rho}}$ variants. a,** *In vitro* autocorrelation traces for 50-150 nM SNAP$^{\text{Rho}}$, AGT54$^{\text{Rho}}$, GE-AGT$^{\text{Rho}}$, ZF5.3-SNAP$^{\text{Rho}}$, ZF5.3-AGT54$^{\text{Rho}}$, and ZF5.3-GE-AGT$^{\text{Rho}}$ were collected in DMEM. All traces were fitted to a 3D diffusion model (solid dark curve) to obtain in vitro diffusion times ($t_{diff}$) as described in Methods. **b,** Representative *in cellulo* autocorrelation traces for Saos-2 cells treated with 1 µM of SNAP$^{\text{Rho}}$, AGT54$^{\text{Rho}}$, GE-AGT$^{\text{Rho}}$, ZF5.3-SNAP$^{\text{Rho}}$, ZF5.3-AGT54$^{\text{Rho}}$, or ZF5.3-GE-AGT$^{\text{Rho}}$ for 0.5 - 2 h. Autocorrelation traces were fitted using a 3D anomalous diffusion model as described in Methods to obtain *in cellulo* diffusion times. *In cellulo* diffusion times are longer than those *in vitro* due to increased viscosity and impaired diffusion in the cytosol.

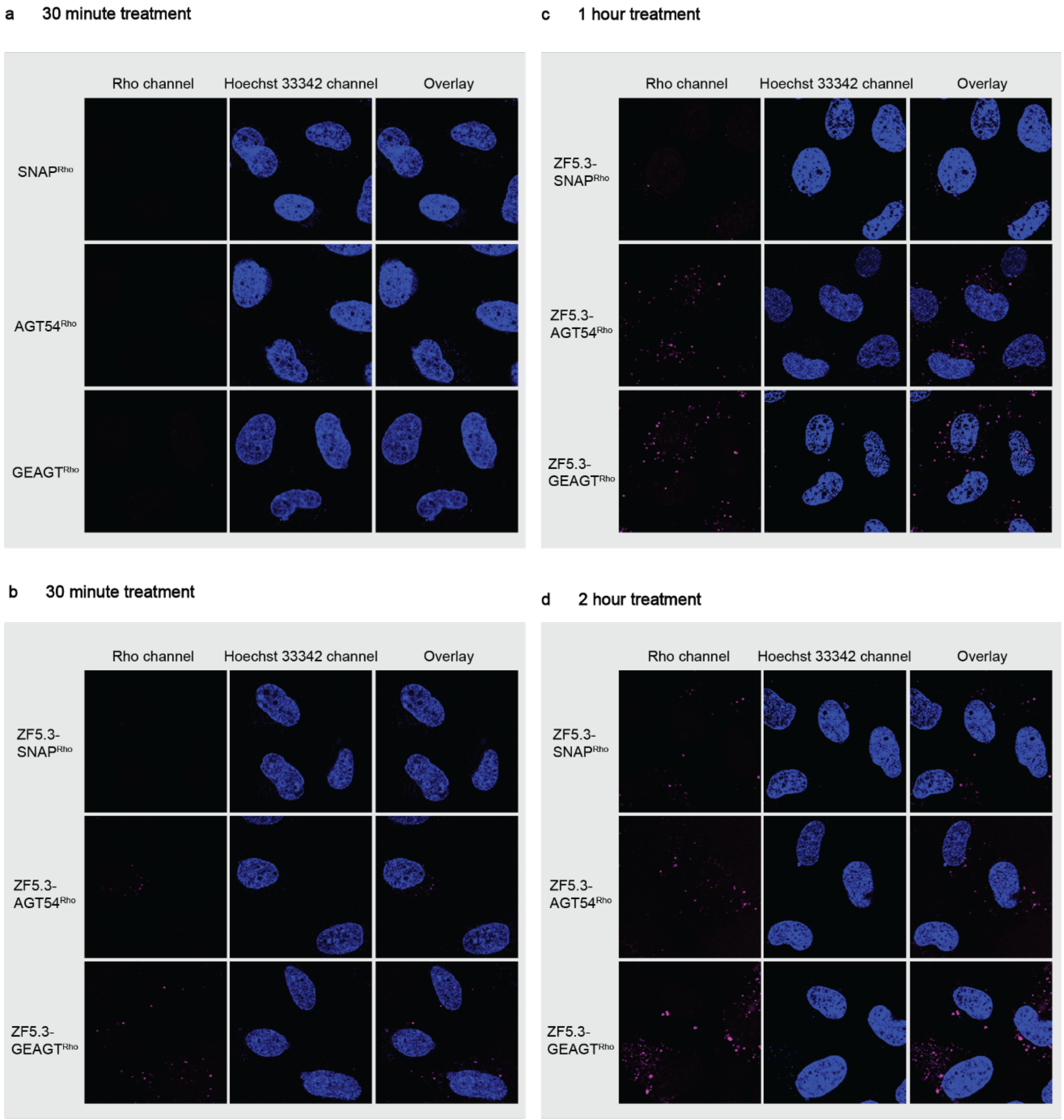


**Fig. S7: 2D confocal microscopy images depicting total intracellular fluorescence of cells treated with SNAP and ZF5.3-SNAP variants.**

**a)** Saos-2 cells were incubated with 1 µM of SNAP$^{Rho}$, AGT54$^{Rho}$, or GE-AGT$^{Rho}$ for 30 min (a) or 1 µM of ZF5.3-SNAP$^{Rho}$, ZF5.3-AGT54$^{Rho}$, or ZF5.3-GE-AGT$^{Rho}$ for 30 min (b), 1 h (c), or 2 h (d), followed by three DPBS washes and trypsinization to remove exogenous protein. Cells were then replated in a fibronectin-coated 8-well microscopy dish and visualized using confocal microscopy. Nuclear fluorescence was detected by adding 300 nM Hoechst 33342 for the final 5 minutes of the protein incubation period. The results shown are representative of at least two biological replicates.

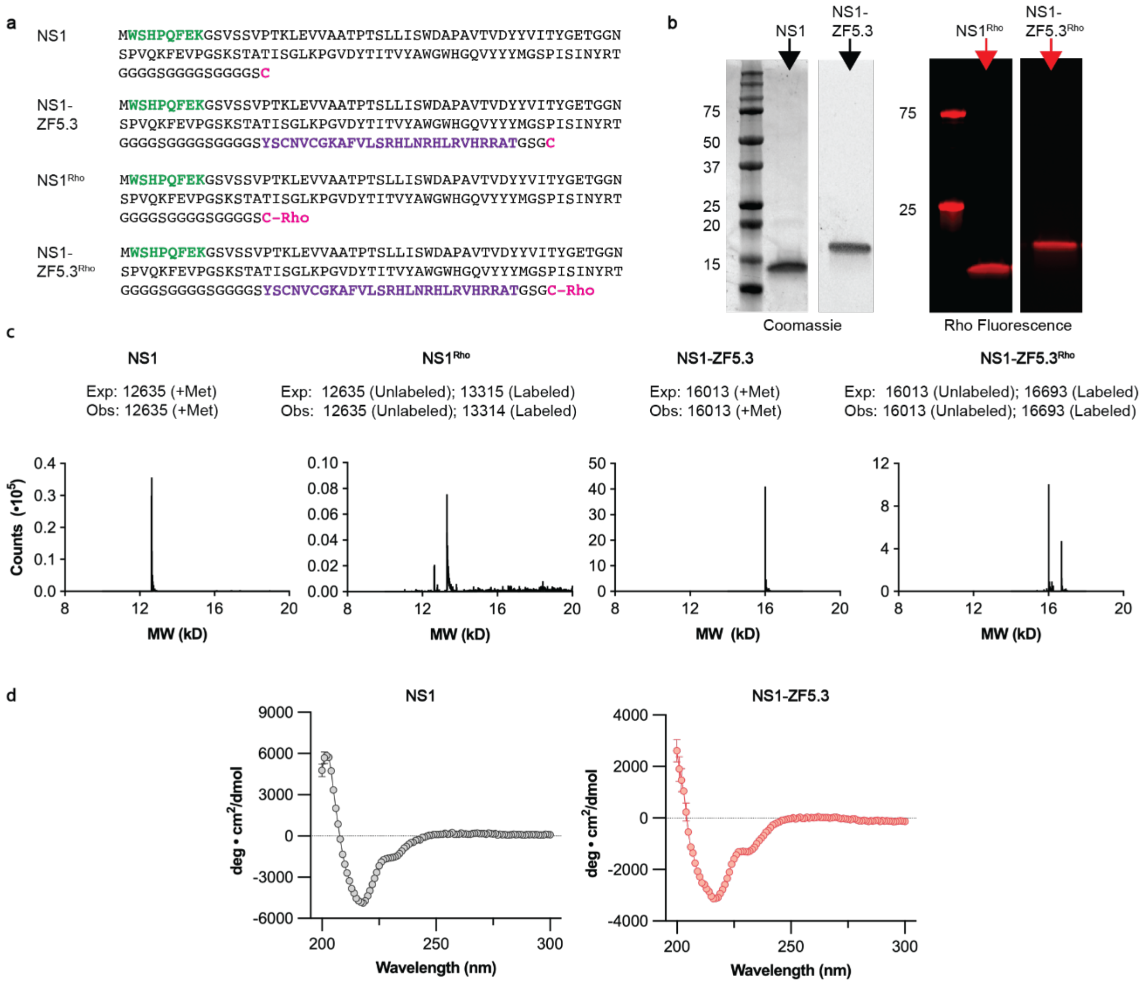


**Fig. S8**: **Purification and characterization of NS1, NS1-ZF5.3, and rhodamine-tagged variants. a,** NS1 proteins chosen for this work included the full-length published NS1 sequence (5) with or without ZF5.3 (purple, 27 aa). All proteins contained an N-terminal Strep-tag for affinity purification (green) and a C-terminal cysteine separated with a flexible linker for conjugation to a maleimide-linked rhodamine dye (magenta). ZF5.3 was appended to the C-terminus of NS1 due to multiple unsuccessful purification attempts with ZF5.3 placed at the N-terminus. **b,** SDS-PAGE analysis of purified NS1, NS1-ZF5.3, $NS1^{Rho}$ and NS1-ZF5.$3^{Rho}$, with proteins visualized either using Coomassie stain (for unlabeled proteins) or fluorescence imaging (for rhodamine-labeled proteins). Labeling efficiency ranged between 20.9 and 50.0%. **c,** LC/MS analysis confirmed the identity of NS1, NS1-ZF5.3, $NS1^{Rho}$, and NS1-ZF5.$3^{Rho}$. For $NS1^{Rho}$ and NS1-ZF5.$3^{Rho}$, prominent peaks were observed for both the unlabeled and rhodamine-labeled species. **d,** Wavelength-dependent CD spectra reveal no significant differences between the secondary structures of NS1 and NS1-ZF5.3. NS1 and NS1-ZF5.3 were measured at 18 µM in 20 mM Tris, 150 mM KCl, 0.5 mM TCEP pH 7.5. Measurements were recorded between 200 and 300 nm in 1 nm intervals with an averaging time of 5 seconds. Data shown are from two biological replicates and are represented as mean ± SEM.

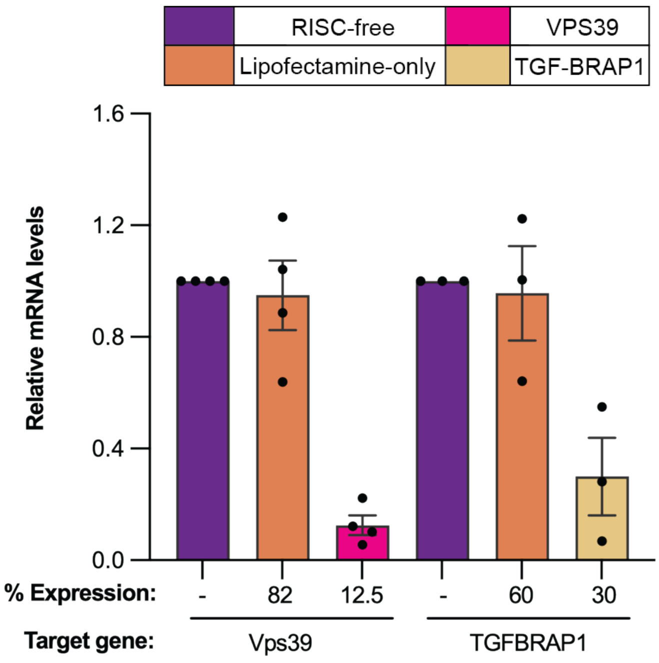


**Fig. S9: qPCR analysis of siRNA knockdowns.** The gene expression level of VPS39 (HOPS complex subunit) and TGF-BRAP1 (CORVET complex subunit) was determined using RT-qPCR and compared to a RISC-free and lipofectamine-only negative control. Data are represented as mean ± SEM with three biological replicates.

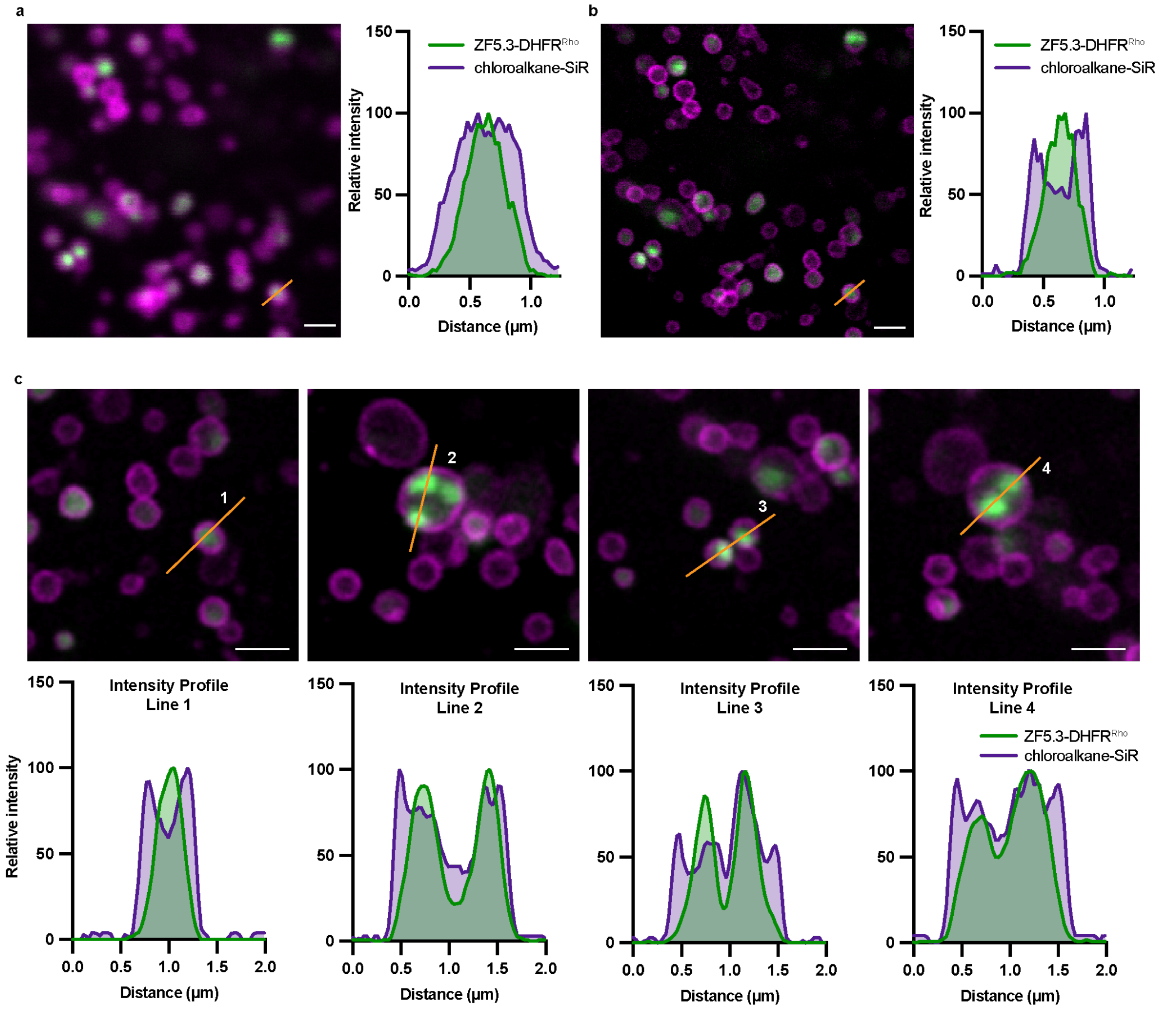


**Fig. S10: ZF5.3-DHFR$^{Rho}$ localizes to the lumen of Lamp1+ vesicles.** Saos-2 cells expressing a Lamp1-HaloTag construct tagged with chloroalkane-SiR were incubated with 0.5 µM ZF5.3-DHFR$^{Rho}$ for 1 hour. Cells were then washed 3x with DPBS, trypsinized to remove exogenously bound protein, and replated in a fibronectin-coated 4-well microscopy dish. **a,** Representative live-cell confocal microscopy images of Saos-2 cells. **b,** TauSTED images depicting lumenal ZF5.3-DHFR$^{Rho}$ within Lamp1+ vesicles. Representative fluorescence intensity line profile shows the relative position of emission from ZF5.3-DHFR$^{Rho}$ (green) and Lamp1-HaloTag-SiR (purple). **c,** TauSTED images depicting sublocalization of ZF5.3-DHFRRho within Lamp1+ vesicles labelled with chloroalkane-SiR. Intensity profiles of lines in each images are graphed below to depict localization of signal from each channel. Scale bars 1 µm.

**Supplementary Tables**

**Table S1: Relevant protein and DNA/RNA Sequences**

| **Protein Sequences** | |
|---|---|
| DHFR | VRPLNCIVAVSQNMGIGKNGDLPWPPLRNEFKYFQRMTTTSSVEGKQNL VIMGRKTWFSIPEKNRPLKDRINIVLSRELKEPPRGAHFLAKSLDDALRLIE QPELASKVDMVWIVGGSSVYQEAMNQPGHLRLFVTRIMQEFESDTFFPEI DLGKYKLLPEYPGVLSEVQEEKGIKYKFEVYEKKLPETGGHHHHHH |
| ZF5.3-DHFR | YSCNVCGKAFVLSRHLNRHLRVHRRATGSGVRPLNCIVAVSQNMGIGKN GDLPWPPLRNEFKYFQRMTTTSSVEGKQNLVIMGRKTWFSIPEKNRPLK DRINIVLSRELKEPPRGAHFLAKSLDDALRLIEQPELASKVDMVWIVGGSS VYQEAMNQPGHLRLFVTRIMQEFESDTFFPEIDLGKYKLLPEYPGVLSEV QEEKGIKYKFEVYEKKLPETGGHHHHHH |
| SNAP-tag | DKDCEMKRTTLDSPLGKLELSGCEQGLHEIKLLGKGTSAADAVEVPAPAA VLGGPEPLMQATAWLNAYFHQPEAIEEFPVPALHHPVFQQESFTRQVLW KLLKVVKFGEVISYQQLAALAGNPAATAAVKTALSGNPVPILIPCHRVVSSS GAVGGYEGGLAVKEWLLAHEGHRLGKPGLGHHHHHH |
| ZF5.3-SNAP-tag | YSCNVCGKAFVLSRHLNRHLRVHRRATDKDCEMKRTTLDSPLGKLELSG CEQGLHEIKLLGKGTSAADAVEVPAPAAVLGGPEPLMQATAWLNAYFHQ PEAIEEFPVPALHHPVFQQESFTRQVLWKLLKVVKFGEVISYQQLAALAGN PAATAAVKTALSGNPVPILIPCHRVVSSSGAVGGYEGGLAVKEWLLAHEG HRLGKPGLGHHHHHH |
| AGT54 | DKDCEMKRTTLDSPLGKLELSGCEQGLHEIKLLGKGTSAADAVEVPAPAA VLGGPEPLMQCTAWLNAYFHQPEAIEEFPVPALHHPVFQQESFTRQVLW KLLKVVKFGEVISYQQLAALAGNPKAARAVKTALSGNPVPILIPCHRVVCS SGAVGGYEGGLAVKEWLLAHEGHRLGKPGLGHHHHHH |
| ZF5.3-AGT54 | YSCNVCGKAFVLSRHLNRHLRVHRRATGSGDKDCEMKRTTLDSPLGKLE LSGCEQGLHEIKLLGKGTSAADAVEVPAPAAVLGGPEPLMQCTAWLNAY FHQPEAIEEFPVPALHHPVFQQESFTRQVLWKLLKVVKFGEVISYQQLAAL AGNPKAARAVKTALSGNPVPILIPCHRVVCSSGAVGGYEGGLAVKEWLLA HEGHRLGKPGLGHHHHHH |
| GE-AGT | DKDCEMKRTTLDSPLGKLELSGCEQGLHEIKLLGKGTSAADAVEVPAPAA VLGGPEPLMQCTAWLNAYFHQPEAIEEFPVPALHHPVFQQESFTRQVLW KLLKVVKFGEVISYQQLAALAGNPKAARAVGGAMRGNPVPILIPCHRVVC SSGAVGGYEGGLAVKEWLLAHEGHRLGKPGLGHHHHHH |
| ZF5.3-GE-AGT | YSCNVCGKAFVLSRHLNRHLRVHRRATGSGDKDCEMKRTTLDSPLGKLE LSGCEQGLHEIKLLGKGTSAADAVEVPAPAAVLGGPEPLMQCTAWLNAY FHQPEAIEEFPVPALHHPVFQQESFTRQVLWKLLKVVKFGEVISYQQLAAL AGNPKAARAVGGAMRGNPVPILIPCHRVVCSSGAVGGYEGGLAVKEWLL AHEGHRLGKPGLGHHHHHH |
| NS1 | WSHPQFEKGSVSSVPTKLEVVAATPTSLLISWDAPAVTVDYYVITYGETG GNSPVQKFEVPGSKSTATISGLKPGVDYTITVYAWGWHGQVYYYMGSPI |

| | |
|---|---|
| | SINYRTGGGGSGGGGSGGGGSC |
| NS1-ZF5.3 | WSHPQFEKGSVSSVPTKLEVVAATPTSLLISWDAPAVTVDYYVITYGETG GNSPVQKFEVPGSKSTATISGLKPGVDYTITVYAWGWHGQVYYYMGSPI SINYRTGGGGSGGGGSGGGGSYSCNVCGKAFVLSRHLNRHLRVHRRAT GSGC |
| StrepTagII-SrtA7m-His6 | WSHPQFEKQAKPQIPKDKSKVAGYIEIPDADIKEPVYPGPATREQLNRGV SFAKENQSLDDQNISIAGHTFIDRPNYQFTNLKAAKKGSMVYFKVGNETR KYKMTSIRNVKPTAVEVLDEQKGKDKQLTLITCDDYNEETGVWETRKIFVA TEVKLEHHHHHH |
| His6-SUMO-SrtA7m | GSSHHHHHHGSGLVPRGSASMSDSEVNQEAKPEVKPEVKPETHINLKVS DGSSEIFFKIKKTTPLRRLMEAFAKRQGKEMDSLRFLYDGIRIQADQTPED LDMEDNDIIEAHREQIGGMQAKPQIPKDKSKVAGYIEIPDADIKEPVYPGPA TREQLNRGVSFAKENQSLDDQNISIAGHTFIDRPNYQFTNLKAAKKGSMV YFKVGNETRKYKMTSIRNVKPTAVEVLDEQKGKDKQLTLITCDDYNEETG VWETRKIFVATEVKLE |
| **Primers** | |
| Primer 1 (to linearize pet-32a(+) vector) | ATGTATATCTCCTTCTTAAAGTTAAACAAAATTATT |
| Primer 2 (to linearize pet-32a(+) vector) | TAACAAAGCCCGAAAGGAAG |
| **siRNA used for RNAi** | |
| RISC-free | siGENOME RISC-Free Control siRNA: D-001220-01 (Dharmacon) |
| KIF11 | siGENOME human SMARTpool siRNA KIF11: D-001220-01 (Dharmacon) |
| VPS39 | siGENOME human SMARTpool siRNA VPS39: M-014052-01 (Dharmacon) |
| TGF-BRAP1 | siGENOME human SMARTpool siRNA TGFBRAP1: M-006903-01 (Dharmacon) |
| **Primers for RT-qPCR** | |
| GAPDH | PrimeTime qPCR primers Hs.PT.39a.22214836 (IDT) |
| VPS39 - Forward | AGGGTCTGGCTATTCCTTATCT |
| VPS39 - Reverse | CCTTGCACCTTCTCACAGTATAG |
| TGF-BRAP1 - Forward | CTGACCACTCAGTACATCATCC |

| | |
|---|---|
| TGF-BRAP1 - Reverse | CTCCTGTCTCCCTATCCTCTT |

**Table S2: Protein Purification Buffers**

| Buffer name | Relevant Protein(s) | Composition |
|---|---|---|
| Lysis Buffer 1 | DHFR proteins, S35 StrepTagII-SrtA7m-His6, His6-SUMO-SrtA7m | 25 mM Tris, 400 mM KCl, 5 mM TCEP, 5% glycerol, pH 7.5 |
| Wash Buffer 1 | DHFR proteins, S35 StrepTagII-SrtA7m-His6, His6-SUMO-SrtA7m | 25 mM Tris, 400 mM KCl, 15 mM imidazole, 5 mM TCEP, 5% glycerol, pH 7.5 |
| Wash Buffer 2 | DHFR proteins, S35 StrepTagII-SrtA7m-His6, His6-SUMO-SrtA7m | 25 mM Tris, 400 mM KCl, 5 mM imidazole, 5 mM TCEP, 5% glycerol, pH 7.5 |
| Elution Buffer 1 | DHFR proteins, S35 StrepTagII-SrtA7m-His6, His6-SUMO-SrtA7m | 25 mM Tris, 300 mM KCl, 250 mM imidazole, 15% glycerol, pH 7.5 |
| Storage Buffer 1 | DHFR proteins, S35 StrepTagII-SrtA7m-His6, His6-SUMO-SrtA7m | 25 mM Tris, 150 mM KCl, 1 mM TCEP, 15% glycerol |
| Lysis Buffer 2 | All SNAP variants | 20 mM Tris, 150 mM NaCl, 10% glycerol, pH 8.0 |
| High-Salt Buffer | All SNAP variants | 20 mM Tris, 1M NaCl, 30 mM imidazole, 10% glycerol, pH 8.0 |
| Low-Salt Buffer | All SNAP variants | 20 mM Tris, 150 mM NaCl, 10% glycerol, pH 8.0 |
| Elution Buffer 2 | All SNAP variants | 20 mM Tris, 150 mM NaCl, 250 mM imidazole, 10% glycerol, pH 8.0 |
| Storage Buffer 2 | All SNAP variants | 20 mM Tris, 150 mM NaCl, 5 mM TCEP, 15% glycerol, pH 8.0 |
| Labeling Reaction Buffer | All SNAP variants | 20 mM Tris, 150 mM NaCl, 10% glycerol pH 7.5 |
| Lysis Buffer 3 | NS1 proteins | 50 mM Hepes, 50 mM NaCl, 10% Glycerol, 1 mM TCEP, 3 mM $MgCl_2$, pH 7.5 |

| | | |
|---|---|---|
| Storage Buffer 3 | NS1 proteins | 50 mM Hepes, 150 mM NaCl, 10% Glycerol, 1 mM TCEP, pH 7.5 |

**Table S3: Sample sizes for all intracellular delivery experiments**

For flow cytometry (FC) experiments, *n* refers to the number of biological replicates. For each biological replicate, 10,000 cells were measured. The "FC Mean" corresponds to the average Median Fluorescence Intensity recorded across all biological replicates. For fluorescence correlation spectroscopy (FCS), *n* refers to the number of individual cell measurements that passed all final filtering criteria (see Online Methods for more details) across all biological replicates. The number of biological replicates for FCS experiments is the same as that shown for the corresponding flow cytometry experiment.

| Condition | Fig | *n* - FC | FC Mean | FC SEM | *n* - FCS | FCS Mean | FCS SEM |
|---|---|---|---|---|---|---|---|
| $DHFR^{Rho}$ - 1 μM, 1h | 1 | 2 | 1085 | 102.9 | 20 | 39.27 | 7.184 |
| $DHFR^{Rho}$ - 0.5 μM, 1h | 1 | 2 | 591.9 | 13.72 | 27 | 34.28 | 4.810 |
| $DHFR^{Rho}$ - 0.1 μM, 1h | 1 | 2 | 304.7 | 7.280 | 35 | 30.38 | 3.483 |
| ZF5.3-$DHFR^{Rho}$ - 1 μM, 1h | 1 | 2 | 33209 | 1073 | 35 | 392.8 | 44.14 |
| ZF5.3-$DHFR^{Rho}$ - 0.5 μM, 1h | 1 | 3 | 9146 | 168.8 | 63 | 348.9 | 24.66 |
| ZF5.3-$DHFR^{Rho}$ - 0.1 μM, 1h | 1 | 2 | 594.1 | 92.30 | 24 | 72.40 | 10.94 |
| $DHFR^{Rho}$ + $ZF5.3^{Unlab}$, 1 μM, 1h | 1 | 2 | 1048 | 4.9 | 21 | 28.42 | 7.518 |
| $DHFR^{Rho}$ - 0.5 μM, 30min | 1 | 2 | 263.9 | 8.475 | 46 | 12.65 | 1.166 |
| $DHFR^{Rho}$ - 0.5 μM, 2h | 1 | 2 | 4067 | 97.75 | 47 | 51.01 | 4.308 |
| ZF5.3-$DHFR^{Rho}$ - 0.5 μM, 30min | 1 | 2 | 8997 | 116.6 | 34 | 216.7 | 28.83 |
| ZF5.3-$DHFR^{Rho}$ - 0.5 μM, 2h | 1 | 2 | 53813 | 1099 | 51 | 360.3 | 36.27 |
| $DHFR^{Rho}$ - 1 μM, 1h + MTX | 2 | 2 | 485.9 | 4.520 | 38 | 17.01 | 2.052 |
| $DHFR^{Rho}$ - 0.5 μM, 1h + MTX | 2 | 2 | 148.0 | 10.78 | 29 | 10.18 | 0.8816 |
| $DHFR^{Rho}$ - 0.1 μM, 1h + MTX | 2 | 2 | 135.6 | 1.130 | 27 | 7.828 | 1.006 |
| ZF5.3-$DHFR^{Rho}$ - 1 μM, 1h + MTX | 2 | 2 | 28322 | 1548 | 36 | 254.2 | 23.09 |
| ZF5.3-$DHFR^{Rho}$ - 0.5 μM, 1h + MTX | 2 | 2 | 5764 | 303.9 | 35 | 173.2 | 19.65 |
| ZF5.3-$DHFR^{Rho}$ - 0.1 μM, 1h + | 2 | 2 | 275.6 | 2.600 | 32 | 21.42 | 1.652 |

| | | | | | | | |
|---|---|---|---|---|---|---|---|
| MTX | | | | | | | |
| ZF5.3-SNAP$^{Rho}$ + MTX - 1 µM, 30min | 2 | 2 | 470.5 | 13.50 | 17 | 19.46 | 2.473 |
| SNAP$^{Rho}$ - 1 µM, 30min | 3 | 3 | 57.33 | 5.840 | 16 | 8.975 | 1.342 |
| AGT54$^{Rho}$ - 1 µM, 30min | 3 | 2 | 199.5 | 1.500 | 20 | 15.34 | 2.672 |
| GE-AGT$^{Rho}$ - 1 µM, 30min | 3 | 2 | 183.0 | 30.00 | 24 | 16.80 | 2.309 |
| ZF5.3-SNAP$^{Rho}$ - 1 µM, 30min | 3 | 3 | 156.0 | 6.110 | 26 | 9.862 | 1.345 |
| ZF5.3-SNAP$^{Rho}$ - 1 µM, 1h | 3 | 2 | 444.5 | 25.50 | 33 | 23.73 | 2.805 |
| ZF5.3-SNAP$^{Rho}$ - 1 µM, 2h | 3 | 2 | 1297 | 43.00 | 48 | 44.24 | 3.493 |
| ZF5.3-AGT54$^{Rho}$ - 1 µM, 30min | 3 | 4 | 1391 | 117.4 | 26 | 64.33 | 5.2777 |
| ZF5.3-AGT54$^{Rho}$ - 1 µM, 1h | 3 | 3 | 2567 | 65.29 | 37 | 85.89 | 6.815 |
| ZF5.3-AGT54$^{Rho}$ - 1 µM, 2h | 3 | 2 | 4435 | 505.0 | 28 | 139.2 | 16.12 |
| ZF5.3-GE-AGT$^{Rho}$ - 1 µM, 30min | 3 | 2 | 2213 | 134.5 | 41 | 133.3 | 16.01 |
| ZF5.3-GE-AGT$^{Rho}$ - 1 µM, 1h | 3 | 2 | 3927 | 533.5 | 20 | 216.2 | 24.53 |
| ZF5.3-GE-AGT$^{Rho}$ - 1 µM, 2h | 3 | 2 | 6795 | 42.00 | 28 | 399.9 | 62.42 |
| NS1$^{Rho}$ - 1 µM, 1h | 4 | 2 | 266.9 | 89.48 | 28 | 17.32 | 2.760 |
| NS1$^{Rho}$ - 2 µM, 1h | 4 | 2 | 1009 | 388.2 | 45 | 23.68 | 3.281 |
| NS1-ZF5.3$^{Rho}$ - 1 µM, 1h | 4 | 2 | 31450 | 985.5 | 43 | 122.9 | 10.31 |
| NS1-ZF5.3$^{Rho}$ - 2 µM, 1h | 4 | 2 | 63974 | 1735 | 32 | 268.3 | 30.16 |
| DHFR$^{Rho}$ - RISC-free siRNA | 5 | 2 | 1005 | 4341.1 | 23 | 24.56 | 2.947 |
| DHFR$^{Rho}$ - VPS39 siRNA | 5 | 2 | 7072.7 | 171.9 | 21 | 24.02 | 2.823 |
| DHFR$^{Rho}$ - TGF-BRAP1 siRNA | 5 | 2 | 1061 | 362.7 | 35 | 22.60 | 2.886 |
| DHFR$^{Rho}$ + MTX - RISC-free siRNA | 5 | 2 | 393.8 | 86.45 | 30 | 11.04 | 1.932 |
| DHFR$^{Rho}$ + MTX - VPS39 siRNA | 5 | 2 | 322.1 | 70.06 | 20 | 8.993 | 1.472 |
| DHFR$^{Rho}$ + MTX - TGF-BRAP1 siRNA | 5 | 2 | 407.9 | 128.8 | 36 | 9.480 | 2.000 |
| ZF5.3-DHFR$^{Rho}$ - RISC-free | 5 | 2 | 36604 | 3815 | 51 | 288.2 | 26.50 |

| | | | | | | | |
|---|---|---|---|---|---|---|---|
| siRNA | | | | | | | |
| ZF5.3-DHFR$^{Rho}$ - VPS39 siRNA | 5 | 2 | 26052 | 1954 | 26 | 140.3 | 15.96 |
| ZF5.3-DHFR$^{Rho}$ - TGF-BRAP1 siRNA | 5 | 2 | 36336 | 3836 | 41 | 422.2 | 61.72 |
| ZF5.3-DHFR$^{Rho}$ + MTX - RISC-free siRNA | 5 | 2 | 11222 | 688.2 | 40 | 209.0 | 14.38 |
| ZF5.3-DHFR$^{Rho}$ + MTX - VPS39 siRNA | 5 | 2 | 8580 | 11.10 | 25 | 177.8 | 22.16 |
| ZF5.3-DHFR$^{Rho}$ + MTX - TGF-BRAP1 siRNA | 5 | 2 | 10689 | 1010 | 38 | 183.5 | 16.11 |
| ZF5.3-SNAP$^{Rho}$ - RISC-free siRNA | 5 | 2 | 1484 | 195.5 | 20 | 55.83 | 7.835 |
| ZF5.3-SNAP$^{Rho}$ - VPS39 siRNA | 5 | 3 | 1174 | 92.93 | 20 | 44.20 | 5.406 |
| ZF5.3-SNAP$^{Rho}$ - TGF-BRAP1 siRNA | 5 | 2 | 2118 | 724.0 | 30 | 51.02 | 4.440 |
| ZF5.3-AGT54$^{Rho}$ - RISC-free siRNA | 5 | 3 | 6258 | 693.6 | 45 | 129.5 | 18.84 |
| ZF5.3-AGT54$^{Rho}$ - VPS39 siRNA | 5 | 4 | 4113 | 371.3 | 15 | 107.3 | 17.15 |
| ZF5.3-AGT54$^{Rho}$ - TGF-BRAP1 siRNA | 5 | 3 | 9150 | 561.9 | 28 | 124.3 | 11.99 |
| ZF5.3-GE-AGT$^{Rho}$ - RISC-free siRNA | 5 | 2 | 5530 | 194.0 | 44 | 291.6 | 30.06 |
| ZF5.3-GE-AGT$^{Rho}$ - VPS39 siRNA | 5 | 2 | 3456 | 348.0 | 28 | 141.6 | 24.40 |
| ZF5.3-GE-AGT$^{Rho}$ - TGF-BRAP1 siRNA | 5 | 2 | 6284 | 83.50 | 42 | 217.1 | 129.5 |
| NS1-ZF5.3$^{Rho}$ - RISC-free siRNA | 5 | 2 | 17505 | 978.5 | 49 | 155.0 | 12.98 |
| NS1-ZF5.3$^{Rho}$ - VPS39 siRNA | 5 | 2 | 14193 | 863.0 | 25 | 99.46 | 12.22 |
| NS1-ZF5.3$^{Rho}$ - TGF-BRAP1 siRNA | 5d | 2 | 18519 | 1778 | 37 | 98.68 | 9.467 |

**SI References**


1. R. F. Wissner, A. Steinauer, S. L. Knox, A. D. Thompson, A. Schepartz, Fluorescence Correlation Spectroscopy Reveals Efficient Cytosolic Delivery of Protein Cargo by Cell-Permeant Miniature Proteins. *ACS Cent. Sci.* **4**, 1379–1393 (2018).
2. S. L. Knox, R. Wissner, S. Piszkiewicz, A. Schepartz, Cytosolic Delivery of Argininosuccinate Synthetase Using a Cell-Permeant Miniature Protein. *ACS Cent. Sci.* **7**, 641–649 (2021).
3. X. Zhang, *et al.*, Dose-Dependent Nuclear Delivery and Transcriptional Repression with a Cell-Penetrant MeCP2. *ACS Cent. Sci.* **9**, 277–288 (2023).
4. B. Mollwitz, *et al.*, Directed Evolution of the Suicide Protein O6-Alkylguanine-DNA Alkyltransferase for Increased Reactivity Results in an Alkylated Protein with Exceptional Stability. *Biochemistry* **51**, 986–994 (2012).
5. R. Spencer-Smith, *et al.*, Inhibition of RAS function through targeting an allosteric regulatory site. *Nat. Chem. Biol.* **13**, 62–68 (2017).
6. Q. Wu, H. L. Ploegh, M. C. Truttmann, Hepta-Mutant Staphylococcus aureus Sortase A (SrtA7m) as a Tool for in Vivo Protein Labeling in Caenorhabditis elegans. *ACS Chem. Biol.* **12**, 664–673 (2017).
7. P. T. Wingfield, Protein Precipitation Using Ammonium Sulfate. *Curr. Protoc. Protein Sci. Editor. Board John E Coligan Al* **APPENDIX 3**, Appendix-3F (2001).
8. N. J. Greenfield, Using circular dichroism spectra to estimate protein secondary structure. *Nat. Protoc.* **1**, 2876–2890 (2006).
9. S. L. Knox, *et al.*, "Chapter Twenty-One - Quantification of protein delivery in live cells using fluorescence correlation spectroscopy" in *Methods in Enzymology*, Chemical Tools for Imaging, Manipulating, and Tracking Biological Systems: Diverse Chemical, Optical and Bioorthogonal Methods., D. M. Chenoweth, Ed. (Academic Press, 2020), pp. 477–505.
10. A. Steinauer, *et al.*, HOPS-dependent endosomal fusion required for efficient cytosolic delivery of therapeutic peptides and small proteins. *Proc. Natl. Acad. Sci.* **116**, 512–521 (2019).
11. J. R. LaRochelle, G. B. Cobb, A. Steinauer, E. Rhoades, A. Schepartz, Fluorescence Correlation Spectroscopy Reveals Highly Efficient Cytosolic Delivery of Certain Penta-Arg Proteins and Stapled Peptides. *J. Am. Chem. Soc.* **137**, 2536–2541 (2015).
12. S. Zheng, *et al.*, Long-term super-resolution inner mitochondrial membrane imaging with a lipid probe. *Nat. Chem. Biol.*, 1–10 (2023).
